# IQI: Individualized Quantitative trait loci (QTL) Index

**DOI:** 10.1101/2025.02.06.636780

**Authors:** Jing Xu, Chen Sun, Junxian Tao, Siyu Wei, Yu Dong, Xuying Guo, Hongsheng Tian, Haiyan Chen, Yingnan Ma, Wenhua Lyu, Shuo Bi, Yuping Zou, Wei She, Chen Zhang, Jingxuan Kang, Fanwu Kong, Mingming Zhang, Hongchao Lyu, Yongshuai Jiang

## Abstract

There is currently no standard efficient method to precisely detecting and characterizing the difference of quantitative trait loci (QTL) effect among groups and/or populations at individual level. The main reason is that the traditional global regression based QTL analyses can only derive an overall effect of genetic variants on the quantitative trait in a specific group. It is a big challenge to obtain the individual-level QTL effect, which is necessary for accurate testing and precise characterization. Here, we first proposed an individualized QTL index (IQI) that represents the degree of regulation between genetic variants and the trait for each individual. Based on IQI, Differential QTL (DQ) analysis framework was presented to accurately detect the case-control differences in more detail. In addition, IQI can be a new feature/biomarker applied to various accurate bioinformatics analysis, such as differential analysis, disease prediction, and survival analysis. Overall, IQI affords a new perspective on pathogenic mechanisms and will facilitate the study of genetic regulation more detailedly and precisely.

## Introduction

Quantitative trait loci (QTL) analysis is an imperative method to understand the association of genetic variation (usually SNP, Single Nucleotide Polymorphism) and the quantitative trait (like height, weight, DNA methylation, gene expression, etc.). With the advance of high-throughput technologies, multiple types of omics data have been generated to facilitate the development of various QTL analysis, such as expression QTL (eQTL), methylation QTL (meQTL), proteomic QTL (pQTL), splicing QTL (sQTL). The QTL analysis has been widely used to elucidate the regulatory mechanism of risk genetic variations and achieved great successes, such as in autoimmune disease [1-5], neurodegenerative disease [6-10], cancer [11, 12].

Of note, the QTL effect may change with individual status [11]. Although it has been aware, the interpretation of QTL results is still limited to simple comparisons of overall global QTL effect between groups. For example, Jennifer A. Webster et al. indicated differences in the regulatory relationship between Alzheimer’s Disease patients and the healthy was only the change of directions (the eQTL effect of rs26133 on the expression level of *AP3M1* was positive in cases but negative in controls) [13]. Kreitmaier et al. identified differential meQTLs potentially with disease stage through comparing the QTL effect existence and directions in low-grade and high-grade osteoarthritis cartilage [14]. It is obviously not accurate, objective, intuitive, comprehensive, and interpretable enough.

In fact, there are many other types of QTL effects differences in the disease state, such as regulatory gain, loss, strengthening, weakening. However, these relationships are hard to be characterized because current QTL are mostly based on overall regression analysis of the quantitative trait (dependent variable) and SNP genotype (independent variable). The regression coefficient (beta) is a global indicator, which is only a single value for each case and control group, so it is not suitable to be used to investigate the detailed characterization of regulatory QTLs. However, it is a big challenge to obtain the individual-level QTL effect. To overcome this hurdle, we first calculate and define the individualized QTL index (IQI) that represents the degree of regulation between SNP and the quantitative trait for each individual and then proposed a standard efficient testing method based on IQI, named Differential QTL (DQ) analysis framework. In addition, the IQI can be utilized as a novel biomarker for many disease analyses.

### Individualized QTL index (IQI) at individual level

To overcome the limitation of traditional global QTL effect (usually be regression coefficient β), the IQI was proposed to represent the degree of regulation between SNP and the quantitative trait at individual level. Given a data set of *n* samples in a specific group (such as cases and controls),

IQI of each individual between a SNP-trait pair ( *g*_*i*_, *t*_*i*_) can be described as (Figure 1A):

**Figure 1.**
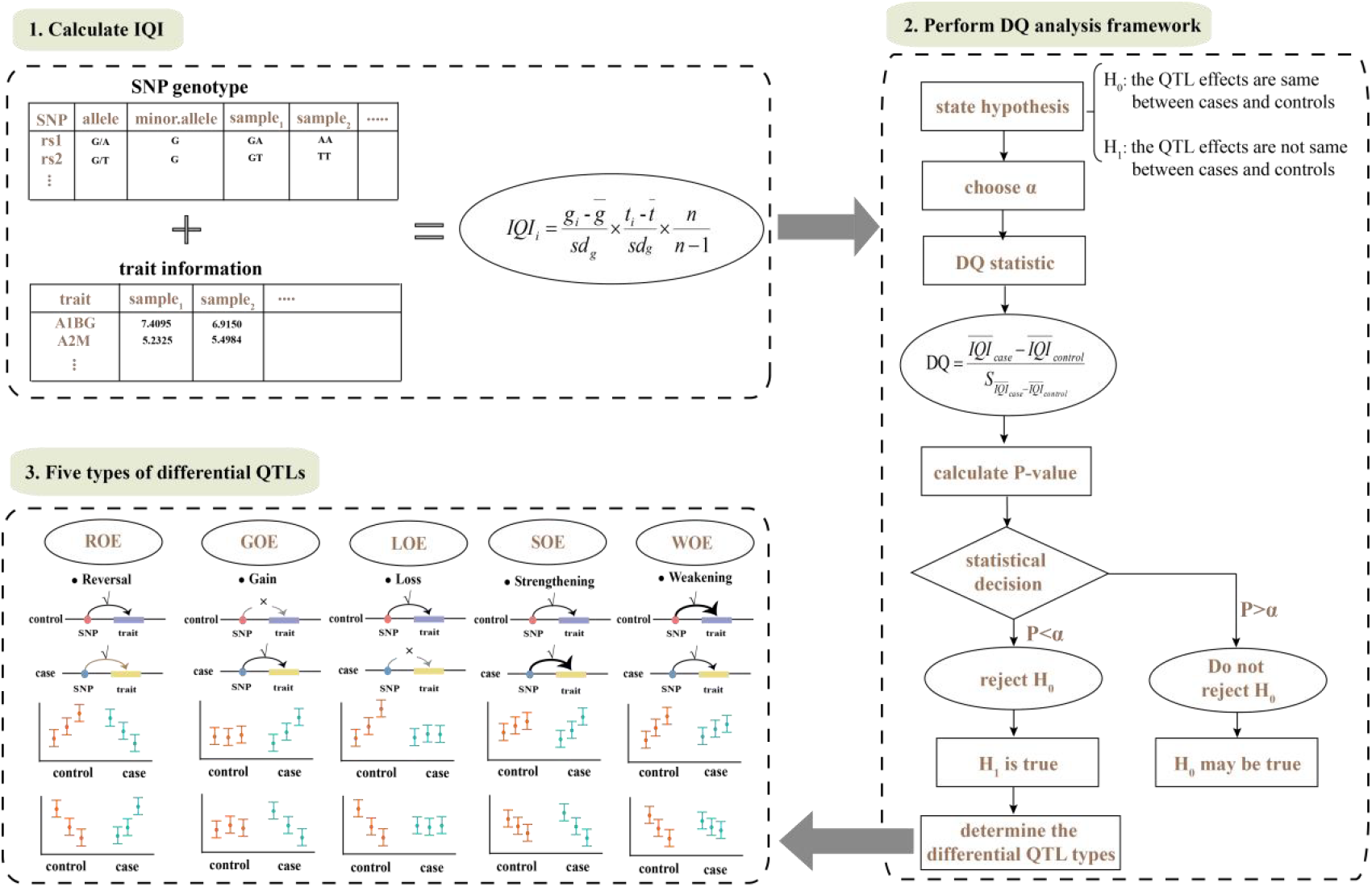
Workflow of IQI calculation and DQ analysis framework. ROE: Reversal of QTL effect; GOE: Gain of QTL effect; LOE: Loss of QTL effect; SOE: Strengthening of QTL effect; WOE: Weakening of QTL effect.

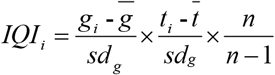

Where *g*_*i*_ denoted the coded genotype (0 for major allele homozygote, 1 for heterozygote, 2 for minor allele homozygote) of sample *i*, and *t*_*i*_ denoted the quantitative trait of sample *i*. *sd*_*g*_ is the standard deviation for coded genotype in the group. *n* is the sample size in the group. The higher the IQI, the stronger the individual regulatory degree.

### The mean IQI is the unbiased estimate of the global regression coefficient β

In fact, the mean IQI was exactly equal to the unbiased estimate of the overall regression coefficient β (Supplementary part 1). Therefore, IQI is a perfect measure of individual QTL regulatory degree. Compared with the global regression coefficient β, IQI provided regulation details at individual resolution, addressing the limitation of traditional approaches that may overlook the individual variation within a group.

### IQI based differential QTL (DQ) analysis framework

IQI at an individual level provided not only a comprehensive view of QTL differences between cases and controls, but also a new solution to accurately and objectively detect the differential QTLs. Based on IQI, we establish the “IQI based differential QTL (DQ) analysis framework” for characterizing QTL effect differences between cases and controls following the steps (Figure 1B):

1. A null hypothesis *H*_*0*_: assuming that the QTL effects (mean IQI) are the same between cases and controls. An alternative hypothesis *H*_*1*_: assuming that the QTL effects (mean IQI) are different between cases and controls.
2. Determined a significance level α and rejection threshold.
3. The test statistics DQ (Difference between QTL effects) was then calculated as

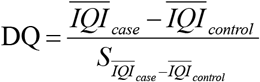 Where 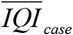 and 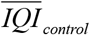 respectively represented the mean IQI for all cases and controls, 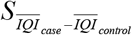 represented the standard error for the difference of mean IQI in cases and controls. was calculated as

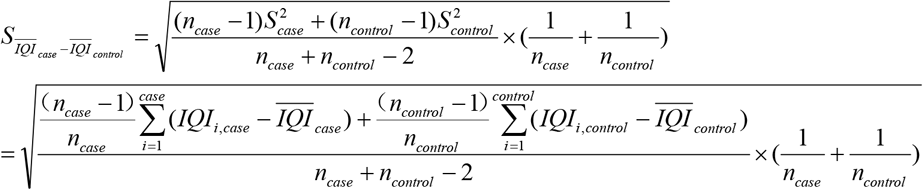

of which, *IQI_i,case_* and *IQI_i,control_* respectively represented the IQI for sample i in case and control group, *n_case_* and *n_control_* respectively represented the sample numbers of case and control group, 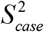 and 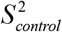 respectively represented the variations of IQI in case and control group. The DQ statistics obey T distribution when with a small sample size and normal distributions with a large sample size (Supplementary part 5).
4. Computed the P value according to the background distribution of DQ statistics.
5. Made a decision. If the P < α, reject the null hypothesis *H*_*0*_ and accept the alternative hypothesis *H*_*1*_, concluding that QTL effects are significantly different between cases and controls.

### Five types of differential QTLs

According to “IQI based DQ analysis framework”, regulatory effect differences can be more accurately and intuitively detected only in one-step. By applying “IQI based DQ analysis framework”, significant differential QTLs could be screened and mainly fall into five types (Figure 1C; Figure 2; Supplementary part 2):

**Figure 2.**
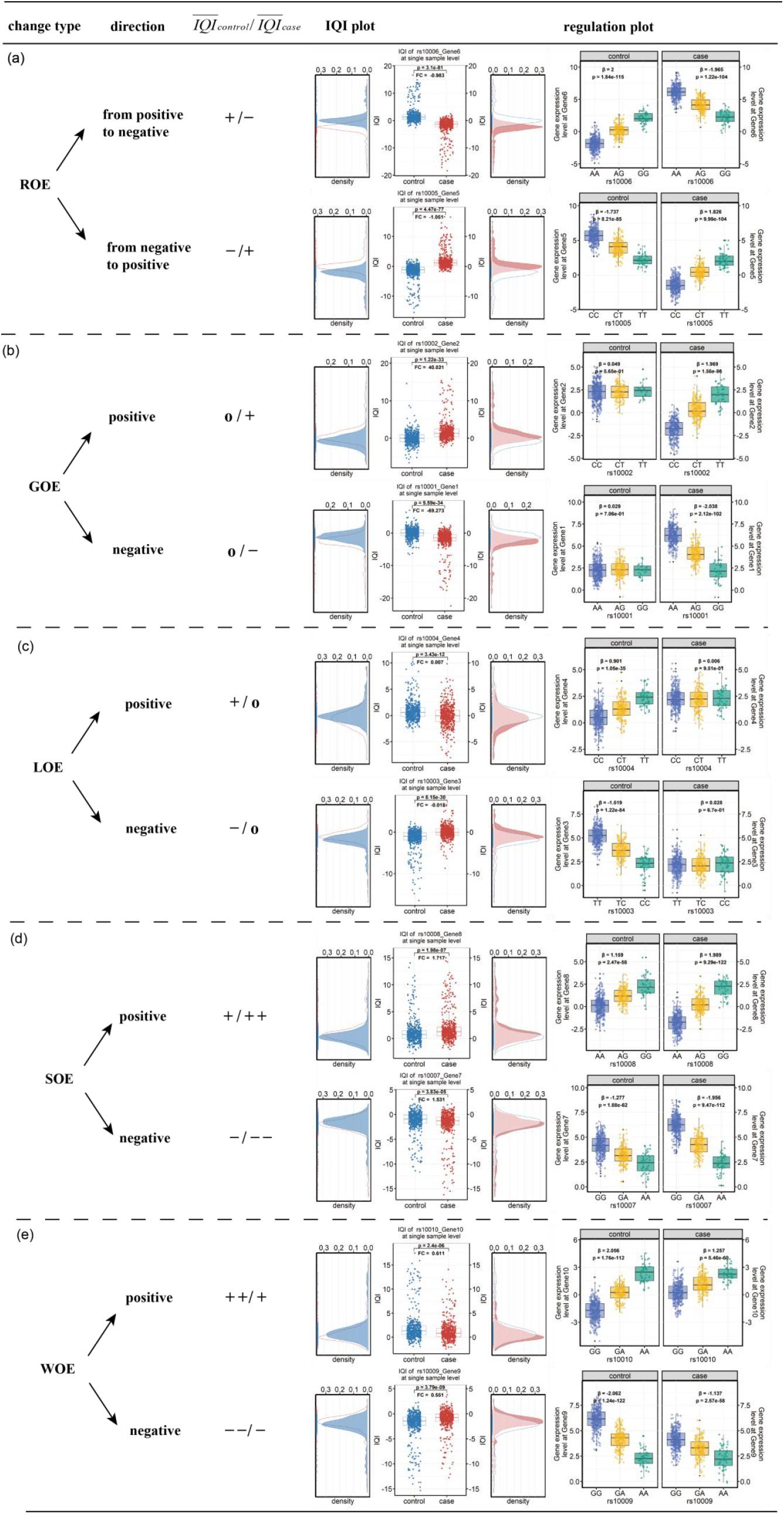
Five types of change regulation generated with simulated data (see more details in Supplementary part 2). +: positive regulation; −: negative regulation; O: insignificant regulation; ++: strengthened positive regulation; --: strengthened negative regulation.

*ROE (Reversal of QTL effect)*: the QTL effects significantly exits both in the normal and disease state but are in an opposite direction. (Figure 2 (a))

*GOE (Gain of QTL effect)*: the QTL regulation only significantly exists in the disease state but not in the normal state. (Figure 2 (b))

*LOE (Loss of QTL effect):* the QTL regulation only significantly exists in the normal state but not in the disease state. In addition, the significant QTL effect can be further divided to positive and negative regulation. (Figure 2 (c))

*SOE (Strengthening of QTL effect)*: the QTL effects significantly exits both in the normal and disease state, but the QTL effect is significantly strengthened in the disease status compared to the normal status. (Figure 2 (d))

*WOE (Weakening of QTL effect)*: the QTL effects significantly exits both in the normal and disease state, but the QTL effect is significantly weakened in the disease state compared to the normal state. (Figure 2 (e))

### IQI as a novel individual feature applying for complex disease analysis

IQI aims to capture and quantify the degree of regulation between genetic variants and quantitative traits at an individual level. In addition to comprehensively detecting QTL differences between case and control groups, IQI offers a unique feature that allows for a more detailed analysis to understand the role of QTL effects in disease pathology. Here, we demonstrated three applications, including differential QTL analysis, disease prediction, and survival analysis.

#### IQI based DQ analysis identified differential QTLs for lung cancer

eQTL analysis provided key insights into the genetic architecture on gene expression levels [15]. It has been conducted in diverse researches, including identifying disease risk loci and/or associated genes, comparing characteristics of populations, analyzing transcriptome data from various tissues and cell types, etc [16, 17]. However, the traditional eQTL analysis was performed in a specific group alone. For example, GTEx is present the largest and most comprehensive eQTL database [18] which only stores eQTL information of 53 non-disease tissues. PancanQTL only stores cancer-related eQTLs [19] that identified using 9,196 tumor samples across 33 cancer type from The Cancer Genome Atlas (TCGA) database. It remains a challenge and barrier to conduct more detailed and accurate analysis of eQTL in different groups. We envision that the “IQI based DQ analysis framework” may surmount this barrier.

Here, we applied “IQI based DQ analysis framework” to 42 lung cancer cases and 42 healthy controls with both genotype and expression information downloaded from the Gene Expression Omnibus (GEO) database under the accession number of GSE33356. After scanning all the regulation pairs of 448,525 SNPs and 24,442 genes, 135 differential eQTLs were identified under the significance level of 2.05e-06 (0.05/24,442). There were 54 GOE (Gain of QTL effect) pairs, 8 LOE (Loss of QTL effect) pairs, and 73 ROE (Reversal of QTL effect) pairs (Figure 3A), involving 127 SNPs and 92 genes on different chromosomes (Figure 4A). Among them, rs4943729_*CT83* was the most significant differential eQTL (p = 1.09e-08) and was divided into GOE type (Figure 4B), which was significant only in lung cancer but not in the healthy state. rs4943729 is located in the intron area of gene *RXFP2* on chromosome 13 with two alleles C and A. *CT83* is on chromosome 11, which has been identified to be associated with diagnosis and prognosis of lung adenocarcinoma [20, 21]. We first reported the QTL difference of rs4943729_*CT83* regulation pair between lung cancer patients and normal controls. With per allele addition of the rs4943729-C, 1.143 addition of gene expression can be observed in lung cancer patients but no significant alteration of gene expression can be observed in healthy controls. The gain of QTL effect of rs4943729_*CT83* in lung cancer patients suggests a potential association that warrants further investigation to elucidate their potential role in the context of lung cancer.

**Figure 3.**
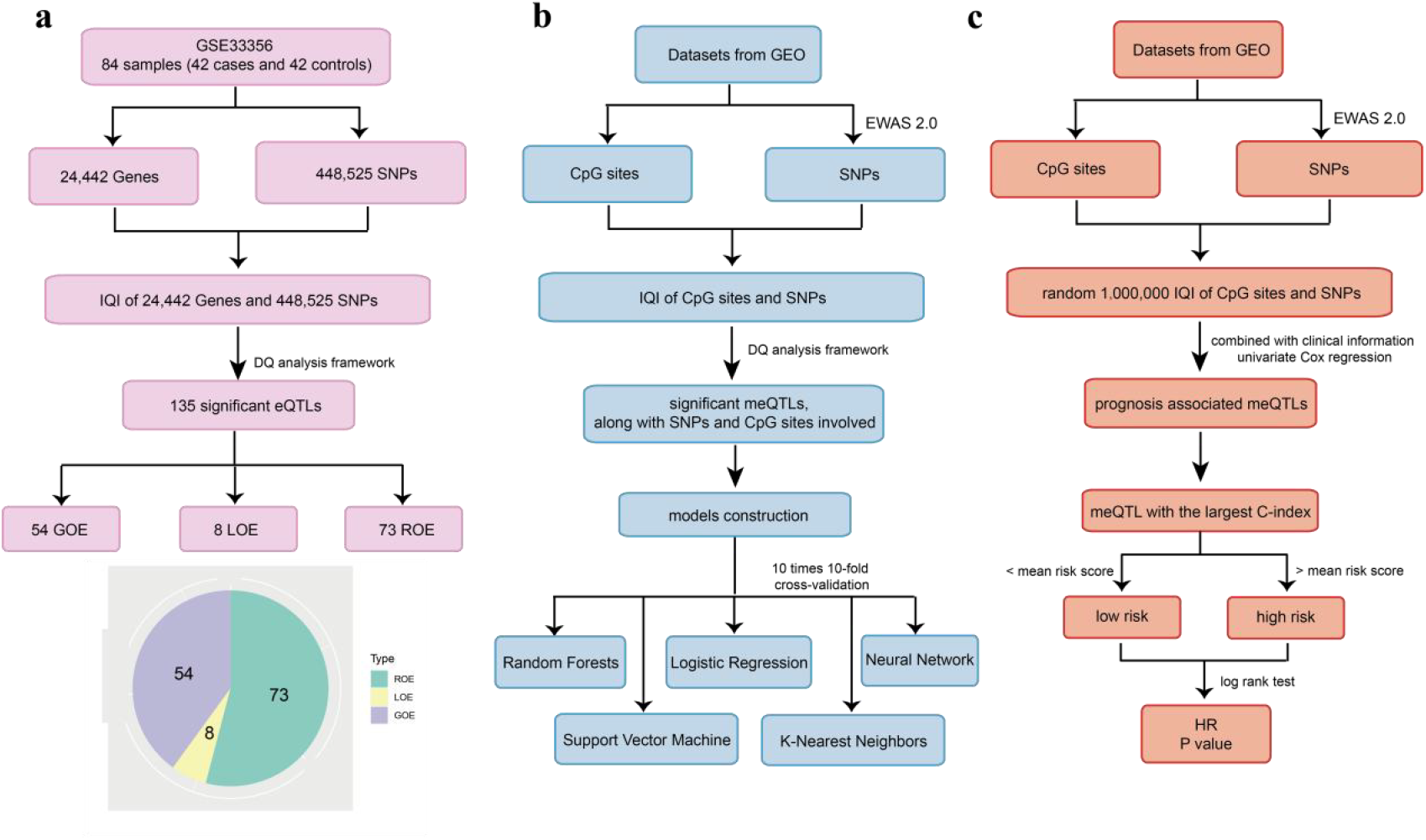
Workflow of three applications that IQI as a novel individual feature for complex disease analysis. (a) IQI based differential QTL analysis identified differential QTLs for lung cancer. (b) IQI based disease prediction. (c) IQI based survival analysis.

**Figure 4.**
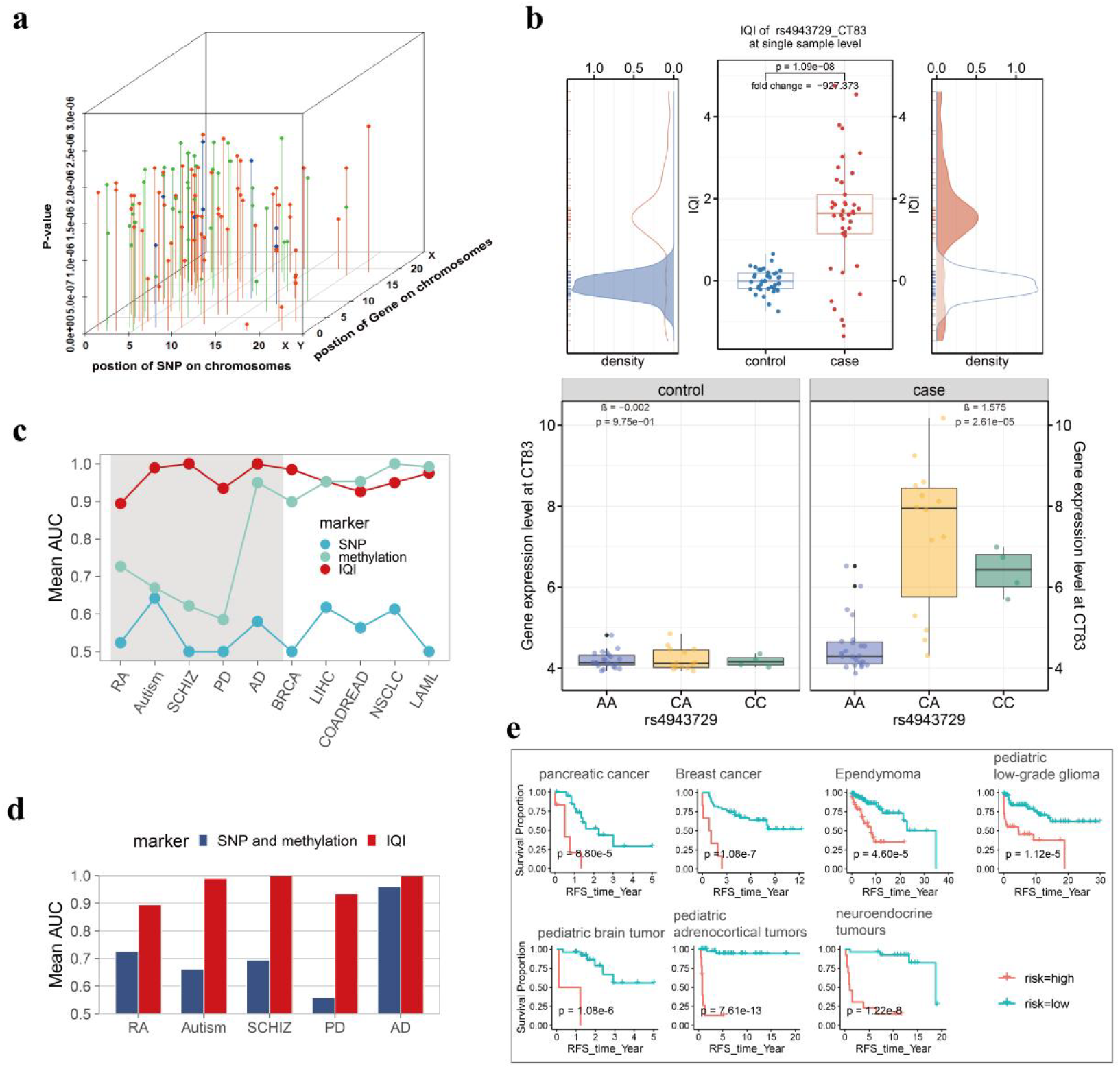
(a) The location of SNP-CpG pairs, the axis-X is the position of SNP, the axis-Y is CpG sites, and the axis-Z is the p value derived by the DQ analysis framework. The red bars are the ROE pairs, blue bars are LOE pairs, and green bars are GOE pairs. (b) The IQI boxplot of rs4943729_*CT83* pair. (c) AUC values of SNP (blue), methylation (green), and IQI (red) for all 10 diseases in SVM model. Gray area in the left half were chronic diseases. (d) AUC values of “SNP and methylation” (dark blue), and IQI (red) for chronic diseases. (e) The Kaplan-Meier (KM) curves of cancer samples. Green and red curves separately indicate low risk and high risk. HR: hazard ratio.

#### IQI based disease prediction

As the degree of regulation between genetic variants and the quantitative trait at individual level, IQI could provide novel insights into how the interaction drives disease susceptibility. It can be employed as a new marker for disease prediction. Here, we comprehensively elucidate the performance and generalizability of IQI through ten prevalent diseases, including five types of chronic diseases (rheumatoid arthritis (RA), Autism, schizophrenia (SCHIZ), Parkinson’s disease (PD), Alzheimer’s disease (AD)) and five types of cancers (breast cancer (BRCA), liver cancer (LIHC), colorectal cancer (COADREAD), non-small cell lung cancer (NSCLC), acute myeloid leukemia (LAML)).

Their DNA methylation data were downloaded from the GEO database (Table S6), covering 485,578 CpG sites. The genotype data of all samples were also derived from the corresponding datasets, involving 62 biallelic SNPs probes designed on the Illumina HumanMethylation450 arrays. All the SNPs used for IQI analysis were ultimately obtained with minor allele frequencies (MAF) > 0.05, missing rate (MR) < 0.01, and Hardy-Weinberg (HWE) p-value > 0.001. This process was performed using the epigenetic association study software EWAS2.0 [22]. To obtain the meQTLs associated with each disease for model construction, we screened all the SNP-CpG pairs by “IQI based DQ analysis framework” (Figure 1) and identified differential meQTLs reaching a secondary significance level of 1e-05 [23]. Subsequently, leveraging IQI of these disease related meQTLs, we developed risk prediction models for each of the diseases utilizing five different machine learning algorithms: Random Forests (RFs), Support Vector Machine (SVM), Neural Network (NN), K-Nearest Neighbors (KNN), and Logistic Regression (LR). To evaluate the performance of these models, we employed a rigorous 10 times 10-fold cross-validation technique (Figure 3B).

SVM and NN models achieved higher performance with more than mean accuracy over 0.93 and mean AUC over 0.94 in the cross-validation across all the ten diseases. RFs and KNN exhibited mean accuracy and AUC over 0.86. In comparison, LR model performed slightly worse with mean accuracy and AUC over 0.78 (Table 1). The results collectively indicated the robust predictive performance of IQI across these diseases. We further tested the performance of SNP, methylation, IQI, and their combined use utilizing SVM model (see more details in Table S7 and Table S8). IQI made substantial predictive performance improvements when the single maker performed poor, especially in chronic diseases (Figure 4C gray area). We observed that IQI improved 37%, 35%, 50%, 43%, 42% AUC beyond the SNP-based model, 17%, 32%, 38%, 35%, 5% AUC beyond methylation-based model, and 17%, 33%, 31%, 38%, 4% AUC beyond ‘SNP+ methylation’-based model for RA, Autism, SCHIZ, PD, and AD prediction (Figure 4C and 4D). Remarkably, even in the cases where single marker already exhibited notable predictive power, IQI incorporated into the model still led to gains in the predictive performance. Although the ‘SNP+methylation’-based model achieved high AUC values of 0.874, 0.946, and 0.948 for BRCA, LIHC, and COADREAD, respectively, the ‘SNP+methylation+IQI’-based model further elevated the model’s performance to AUC values of 0.983, 0.978, and 0.963. Moreover, for NSCLC and LAML, both surpassed AUC values of 0.99 (Table S7 and Table S8).

**Table 1.** The accuracy and AUC of prediction performance with IQI across ten diseases.

| Disease | Mean Accuracy |  |  |  |  | Mean AUC |  |  |  |  |
| --- | --- | --- | --- | --- | --- | --- | --- | --- | --- | --- |
|  | SVM | NN | RFs | LR | KNN | SVM | NN | RFs | LR | KNN |
| RA | 0.895 | 0.870 | 0.853 | 0.820 | 0.815 | 0.894 | 0.870 | 0.853 | 0.820 | 0.816 |
| Autism | 0.990 | 0.978 | 0.854 | 0.911 | 0.961 | 0.990 | 0.978 | 0.854 | 0.910 | 0.962 |
| SCHIZ | 1.000 | 0.986 | 0.919 | 0.721 | 0.990 | 1.000 | 0.985 | 0.899 | 0.721 | 0.989 |
| PD | 0.938 | 0.943 | 0.845 | 0.657 | 0.881 | 0.934 | 0.943 | 0.823 | 0.654 | 0.880 |
| Alzheimer | 0.999 | 0.998 | 0.996 | 0.626 | 1.000 | 1.000 | 0.998 | 0.996 | 0.628 | 1.000 |
| BRCA | 0.984 | 0.940 | 0.864 | 0.904 | 0.869 | 0.985 | 0.947 | 0.581 | 0.887 | 0.922 |
| LIHC | 0.954 | 0.895 | 0.938 | 0.851 | 0.771 | 0.953 | 0.895 | 0.938 | 0.852 | 0.771 |
| COADREAD | 0.926 | 0.894 | 0.983 | 0.612 | 0.742 | 0.926 | 0.894 | 0.983 | 0.612 | 0.742 |
| NSCLC | 0.950 | 0.959 | 0.901 | 0.866 | 0.860 | 0.950 | 0.959 | 0.901 | 0.866 | 0.860 |
| LAML | 0.981 | 0.924 | 0.925 | 0.883 | 0.743 | 0.975 | 0.935 | 0.872 | 0.883 | 0.825 |

In summary, IQI is a novel and efficient disease predictive characteristic. It provides individual information of how specific genetic variants may interact with quantitative traits beyond single markers (such as DNA methylation). Our findings support the notion that the IQI can effectively serve as a valuable indicator for disease prediction and medical decision-making. Importantly, IQI does not conflict with other markers (such as SNPs and methylation). In practical applications, IQI can be combined with other biomarkers to enhance predictive performance.

#### IQI based survival analysis

The prognosis of patients is another one of the primary aspects for complex disease analysis [24-26]. The QTLs have emerged as potential prognostic biomarkers in various human disease. Similarly, IQI of the risk QTLs provide new insight into disease prognosis. Here, we demonstrated the example of introducing IQI of meQTLs as biomarkers for the survival of seven malignant tumor, including pancreatic cancer, breast cancer, ependymoma, pediatric low-grade glioma, pediatric brain tumor, pediatric adrenocortical tumor, and neuroendocrine tumor.

The DNA methylation data for all patients were downloaded from the GEO database (Table S9). The corresponding genotype data designed on Illumina HumanMethylation450 arrays or Infinium MethylationEPIC BeadChip for these samples were obtained using the EWAS2.0 software [22]. Those SNPs used for analysis all passed the same filtering criteria as “IQI based disease prediction”. The clinic information, including the live state and survival time (in years), was obtained from the GEO database or the supplementary file of the original paper [27]. Considering the extensive computational burden, we randomly selected a subset of 1,000,000 SNP-CpG pairs for IQI calculation and subsequent prognosis analysis. By combining the IQI data with the clinical information, the meQTLs associated with prognosis were identified through univariate Cox regression analysis with the R packages ‘survival’ and ‘Hmisc’ under the significance level of 0.001. Further, the meQTLs with highest concordance index (C-index) were used for prognosis prediction. Patients were categorized into low-risk (greater than mean risk score) and high-risk (less than mean risk score) groups according to the predicted risk scores. The log-rank test was utilized to evaluate the difference in overall survival between the different risk groups (Figure 3C).

Results showed that IQI achieved commendable prediction performance across all the tumors, with the highest C-index values were more than 0.74 in ependymoma and pediatric low-grade glioma, 0.81 in breast cancer and neuroendocrine tumor, 0.90 in pancreatic cancer and pediatric brain tumor, and even 0.95 in pediatric adrenocortical tumor. Notably, the overall survival (OS) rate of patients in the low-risk group was significantly higher than that in the high-risk group, showing Hazard ratio (HR) of 6.036 (P-value=8.80e-5, pancreatic cancer), 7.779 (P-value=1.08e-7, breast cancer), 3.558 (P-value=4.60e-5, ependymoma), 3.857 (P-value=1.12e-5, pediatric low-grade glioma), 17.217 (P-value=1.08e-6, pediatric brain tumor), 33.834 (P-value=7.61e-13, pediatric adrenocortical tumor), and 10.443 (P-value=1.22e-8, neuroendocrine tumor), respectively (Figure 4E). In addition, we conducted a comparative analysis of the predictive performance of SNP, methylation, IQI, and the combined use for prognosis risk assessment. The findings revealed that when single markers (such as SNP and methylation) were not enough to be employed as significant prognostic biomarkers, IQI led to substantial improvements in predictive performance (Table S10). Specifically, IQI improved the C-index beyond SNP-based model by 44% (0.510 to 0.950), methylation-based model by 45% (0.504 to 0.950), and ‘SNP+methylation’-based model by 45% (0.496 to 0.950) in pediatric adrenocortical tumor. Remarkably, although the combination of SNP and methylation could achieve a relatively high predictive performance in neuroendocrine tumor, with a C-index of 0.878, the incorporation of IQI into the model further enhanced the predictive performance to 0.913 (Table S10). All the results indicated that IQI can be a robust biomarker for the prognostic prediction, offering valuable insights into disease prognosis.

### Advantages of IQI over traditional linear regression for QTL analysis

Currently, QTL analysis are mainly based on traditional linear regression. However, the regression coefficient is a global indicator which is only a single value for a group sample. IQI is an individual-focused, innovative, and more detailed index representing the degree of regulation for each single sample at the individual level. In addition, the traditional regression based differential QTL identification is just through simple comparisons of the overall regression coefficients between groups, such as size and orientation. IQI based DQ framework enables dividing QTL effect differences into the more detailed five types (ROE, GOE, LOE, SOE, and WOE) directly and precisely. Furthermore, IQI can be applied as a novel individualized indicator for subsequent finer analysis, such as clustering analysis, principal component analysis (PCA), survival analysis, and network analysis, while traditional regression can not.

In summary, IQI offers a clearer and more efficient method to enhance our understanding of genetic interactions and their implications in disease contexts from a novel perspective.

#### Software for IQI calculation and IQI based differential QTL analysis

The IQI calculation and “IQI based DQ analysis framework” were accomplished by the JAVA language and integrated as a software platform, which is free available at http://www.onethird-lab.com/IQI/. The software required four input files including the genetic variation (SNP genotype) data and the quantitative trait information of each sample in cases and controls (Supplementary part 3). Noteworthy, samples should be consistent in order. The final output provided IQI of all samples, DQ analysis results (including DQ statistics, average IQI in cases and controls, Fold-change, change type of QTL effects), and the traditional regression based QTL analysis results. The IQI boxplot also can be viewed.

## Discussion

In this work, we motivate the individualized QTL index (IQI) for the degree of regulation between genetic variants and the quantitative trait for each individual. IQI remedies the limitations that traditional regression coefficient is only a single value for one group and not suitable to be used to compare QTL effect differences among groups. The regression analysis has been used for detecting the relationships between gene-environment interactions and diseases. For example, Biernacka et al. conducted the gene-environment interaction analysis of pesticide exposure and risk of Parkinson’s disease [28]. They first performed the conditional logistic regression to fit models with the binary pesticide exposure variable and the SNP genotype, with and without the interaction term between these two variables, and then applied the one degree-of-freedom likelihood ratio test of the interaction effect to compare the two models [28]. Else, candidate gene analyses, genome-wide interaction and association studies (GWAIS), and genome-wide and exposome-wide association studies (GxEWAS) have also been used frequently [29]. However, there are many limitations. One major limitation is that these methods typically rely on aggregated measures of quantitative traits, such as average levels across a population/group (cases or controls), disregarding the individual-level variations that may exist. Moreover, these methods should be carried out through multiple steps and the interpretation of results is not detailed and accurate.

On the basis of IQI, we performed the DQ analysis framework, which can be applied to detect the QTL effect differences between healthy and disease state. IQI based DQ analysis framework is a novel method achieving the one-step process, efficient, and convenient performance. According to DQ analysis framework, QTL effect differences can be divided into five typical significant types: ROE, GOE, LOE, SOE, and WOE. These QTL differences afford a new perspective on pathogenic mechanisms. In addition, IQI can be a novel and potential biomarker for various accurate and elaborate bioinformatics analysis. We demonstrated the three applications of IQI, including differential QTL analysis, diagnosis analysis, and survival analysis. The results proved that IQI is a strong and robust biomarker for facilitating our understanding of the genetic regulatory effect on the quantitative trait in complex disease. Noteworthy, the example applications were independent to each other and the datasets used were different and disconnected. Each dataset in each application was from the same study, which was not confounded with batch effects. In the special study, if multiple datasets that from different populations or studies were used, the batch effect should be removed first to ensure accurate results.

IQI is the degree of regulation between genetic variants and the quantitative trait at the individual level. It is a novel and efficient indicator that can be employed as a new marker for disease prediction and survival analysis, encompassing machine learning, deep learning, etc. Importantly, IQI does not conflict with other markers. If there is a strong interaction between a SNP and the quantitative trait, the introduce of IQI to models will achieve higher performance. In practical applications, we recommend considering combination of IQI with other markers (such as SNP and methylation) to obtain more comprehensive information.

Furthermore, to further substantiate the meaningfulness of IQI in practical applications and validate that IQI is not simply an unexplained error, we compared the predictive performance for RA utilizing the IQI before and after a complete shuffling (where the regulatory relationships between SNPs and quantitative traits would be disrupted) (Supplementary Part 7, Figure S9). We conducted tests using five models - support vector machines (SVM), neural networks (NN), random forests (RFs), Logistic Regression (LR), and K-Nearest Neighbors (KNN). The ten-fold cross-validation method, along with the assessment metrics of AUC, was employed to evaluate the predictive performance. Theoretically, if IQI is simply an unexplained error, then the predictive performance of IQI should remain relatively unchanged. Intriguingly, we observed that the predictive performance for IQI before the shuffle was significantly higher than that after the shuffle (Figure S10). Furthermore, after the shuffle, the mean AUC of IQI in the five models are all around 0.5 (0.523 for SVM, 0.521 for NN, 0.535 for LR, 0.523 for KNN, and 0.634 for RFs), whereas the mean AUC before the shuffle are all above 0.8 (0.902 for SVM, 0.869 for NN, 0.862 for RFs, 0.824 for LR, and 0.820 for KNN) (Figure S10). This further underscores our interpretation of IQI serving as an individualized QTL indicator rather than simply unexplained random error.

To conclude, our study highlights the need for considering QTL effect at individual-level. We anticipate our approach will facilitate the study about genetic regulatory mechanisms more precisely and its scope will amplify with the increasing availability of the multi-omics data.

## Acknowledgements

This work was supported by the National Natural Science Foundation of China [Grant Nos. 31970651, 92046018]; Mathematical Tianyuan Fund of the National Natural Science Foundation of China [Grant No. 12026414]; Basic Research Support Project for the Excellent Youth Scholars of Education Department of Heilongjiang Province [Grant No. YQJH2023036].

## Data Availability

All data to support our findings are available through http://www.onethird-lab.com/IQI/ and https://github.com/onethird-lab/IQI.

## Author Contributions

J.X., C.S., J.X.T. and S.Y.W. contributed equally to this work. Y.S.J. conceived and contributed the work. J.X., C.S., J.X.T., S.Y.W.,Y.D., X.Y.G., H.S.T, H.Y.C., W.H.L., S.B., Y.P.Z., W.S., C.Z., J.X.K., F.W.K., M.M.Z., H.C.L., and Y.S.J. drafted and modified the manuscript. Y.N.M. are important contributors of the Funome Project. The Funome Project provided data support.

## Supplementary Materials for

### 1. The mean IQI is the unbiased estimate of the global regression coefficient β

#### 1.1 The mean of IQI

IQI of each single sample between an SNP-trait pair ( *g*_*i*_, *t*) can be calculated as:

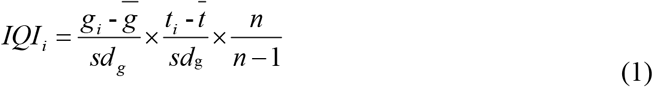

The mean IQI is

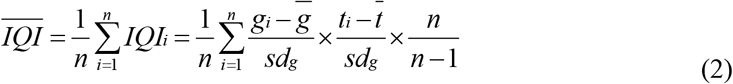

#### 1.2 The regression coefficient β

The traditional regression equation is

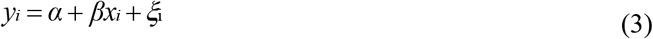

where *α* and *β* are two constants, *ξi* is an unobservable variable. The coefficient β was deduced as:

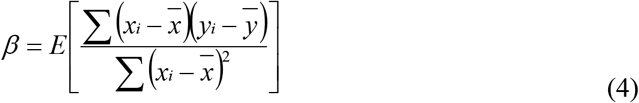

#### 1.3 Proof of that the mean IQI is the unbiased estimate of β

We find that the mean IQI was exactly equal to the unbiased estimate of the overall regression coefficient β, i.e. 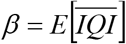

Proof: let *xi* = *gi, yi* = *ti* in the Eq (4), i.e.

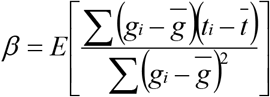

And

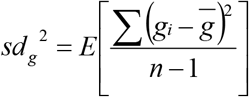

So that

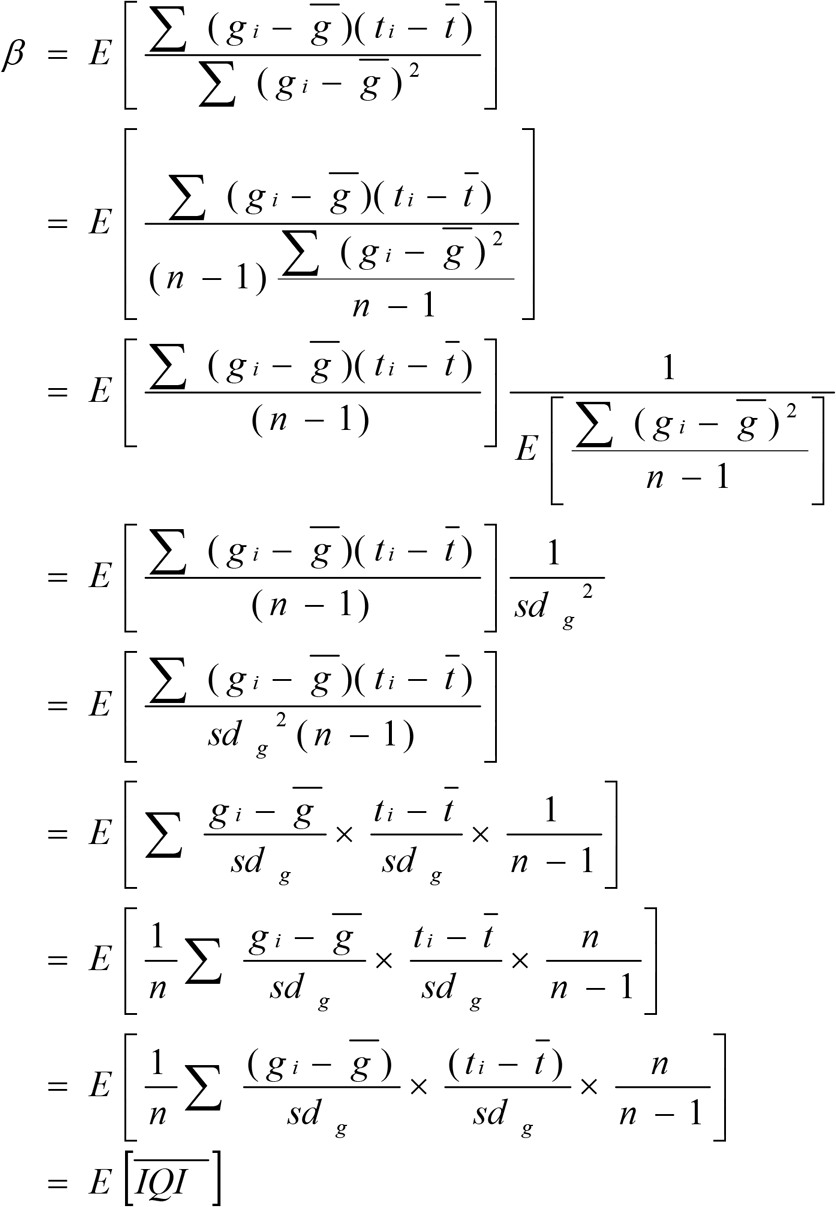

### 2. DQ analysis framework with simulated data to identify differential QTLs

#### 2.1 Simulated data generation

As the scant sample data with both gene expression and SNP genotype, we used the simulated data to perform the DQ analysis framework.

##### Genotype mdata

The genotype data set for case group was generated according to the following rules: for each SNP, (i) the minor allele was defined as C, while the non-minor allele was defined as T, and (ii) the minor allele frequency (MAF) across all samples was set to be at least 0.5.

Similarly, the genotype data set for control group was generated according to the following rules: for each SNP, (i) the minor allele was defined as C, while the non-minor allele was defined as T, and (ii) the minor allele frequency (MAF) across all samples was set to be at least 0.2.

Following these rules, we generated genotype data sets for the case group with a sample size of 500 and 500 SNPs, as well as for the control group with the same sample size and SNP count.

##### Gene expression data

The gene expression data were generated based on the above genotype data according to the interaction regression formula:

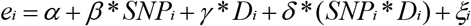

where *e*_*i*_ is gene expression of sample *i*; *α* is the intercept; *β*is the correlation coefficient between SNP and gene expression; *SNP*_*i*_ is the coded genotype (0 for major allele homozygote, 1 for heterozygote, 2 for minor allele homozygote) of sample *i*; *γ*is the correlation coefficient between disease status and gene expression; *D*_*i*_ is the coded disease status (1 in cases and 0 in controls) of sample *i*; *δ* is the correlation coefficient between interaction of SNP and disease status and gene expression; *ξ_i_* is the error term.

During the stimulate data generation, *α* was random number following the normal distribution with mean of 2 and standard deviation of 1; *β*,*γ*, and *δ* were randomly-distributed number from −2 to 2; *ξ_i_* was random number following the normal distribution with mean of 0 and standard deviation of 1.

For the case group, let *D*_*i*_ as 1, then the gene expression data can be obtained according the genotype data of 500 samples in the case group as following:

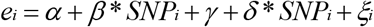

For the control group, let *D*_*i*_ as 0, then the gene expression data can be obtained according the genotype data of 500 samples in the control group as following:

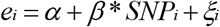

Collectively, we obtained 500 simulated SNPs genotype and genes expression data of 500 cases and 500 controls, respectively. We performed the above process with R language as described below.

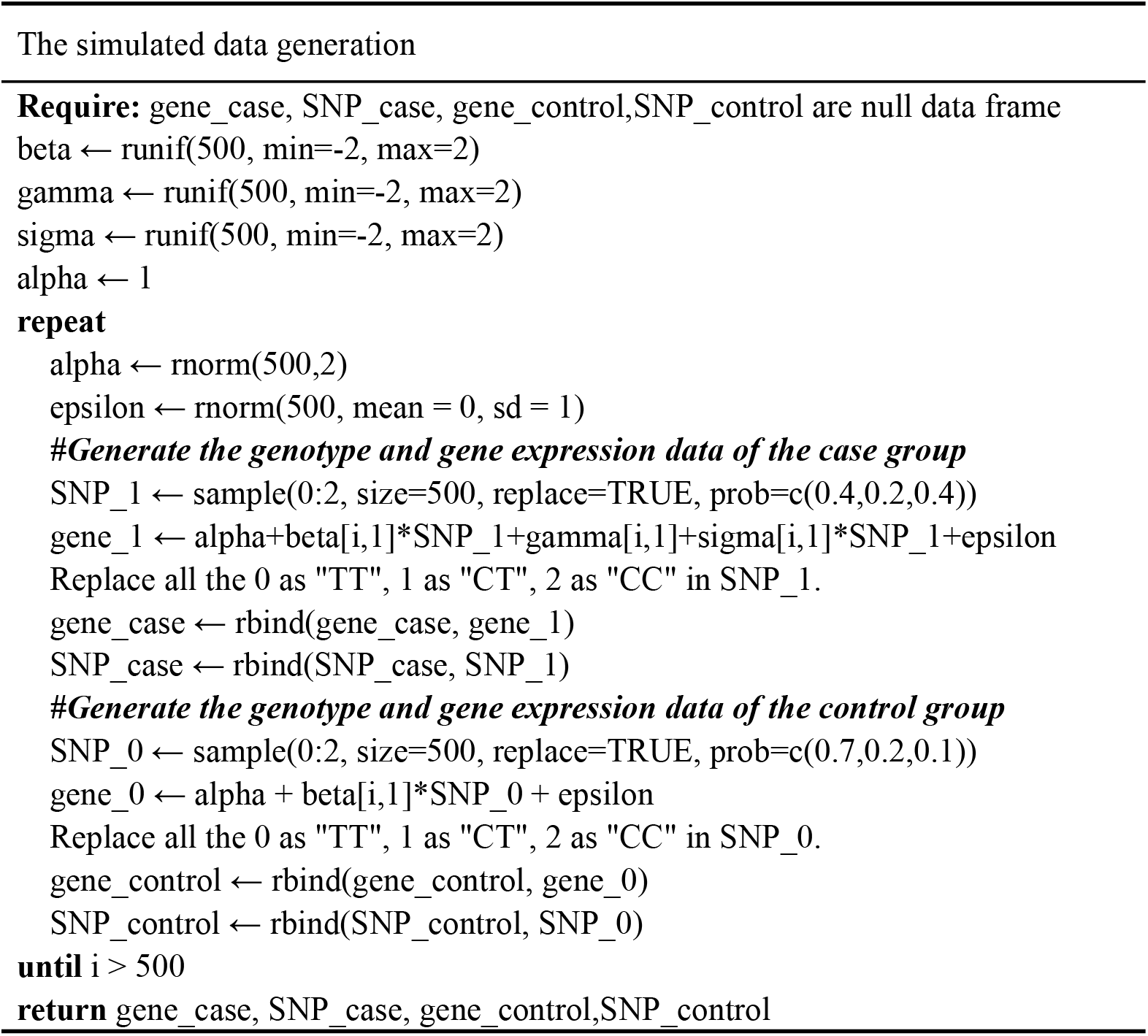

#### 2.2 Determined the differential QTL type

There are mainly five types of significant QTL effect differences, including ROE, (Reversal of QTL effect), GOE (Gain of QTL effect), LOE (Loss of QTL effect), SOE (Strengthening of QTL effect), and WOE (Weakening of QTL effect) type according to the regulation differences between cases and controls. We decide the difference type by comparing the P-values in the cases and controls and the significance level α. For example, if the P-value was less than α in the case but more than α in the control, it will be classified as a GOE type. Each of them can be further divided into two more detailed types (positive or negative regulation) according to the mean IQI in case and control groups and fold change. If the mean IQI is greater 0, it is a positive regulation, otherwise, it is negative. Based on the fold change, the similar regulation can be divided into strengthening or weakening. See the table bellow for detailed information (Table S1).

**Table S1.**
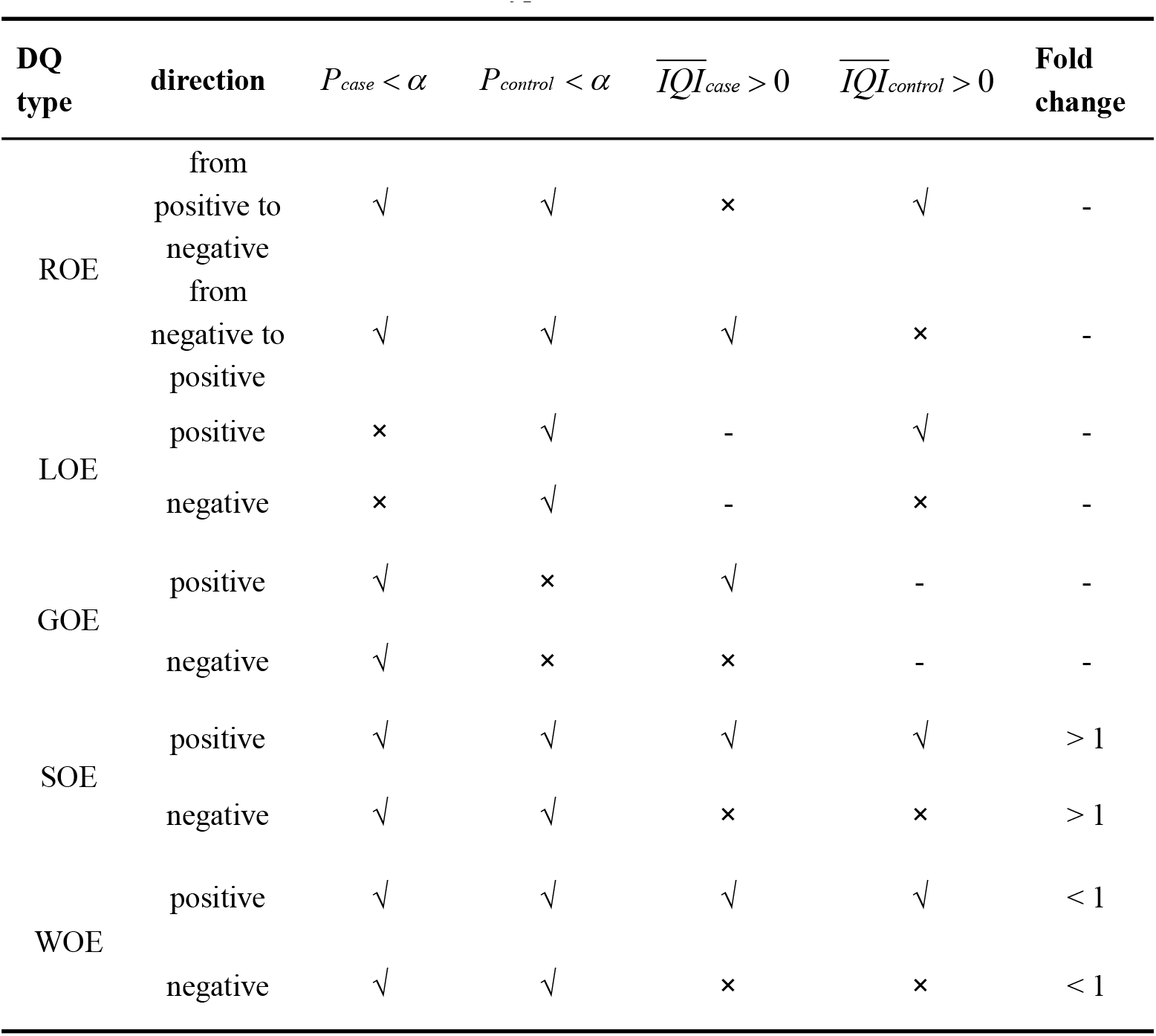
The ten differential QTL (DQ) types.

#### 2.3 The result of DQ analysis with simulated data

The significant regulatory SNP-gene pairs were identified with the significance level of 1e-04 (0.05 divided by the total gene count of 500). After performing the DQ analysis framework, 434 differential eQTLs were identified and five types of QTL effect differences were obtained (35 GOE, 23 LOE, 217 ROE, 101 SOE, and 58 WOE).

The first type was GOE (Gain of QTL effect), where the QTL effects was significant only in the disease state but not in the normal state (Figure S1). The absolute mean IQI was less than 0.05 in the control but more than 0.05 in the case (top half of Figure S1), which was equal to the regression coefficient β (bottom half of Figure S1). In addition, the density of IQI between the cases and controls (both sides of the top half of Figure S1) is significantly different. The GOE type can be further divided into the gain of positive (Figure S1A) and negative regulation (Figure S1B). In accordance with the IQI, the regulatory effect on SNP to gene expression was not significant in the control (p > 0.05) but positive (p < 0.05, the mean IQI > 0, β > 0) in the case (the lower half of Figure S1A). In Figure S1B, the regulatory effect was not significant in the control (p > 0.05) but negative (p < 0.05, the mean IQI < 0, β < 0) in the case.

**Figure S1.**
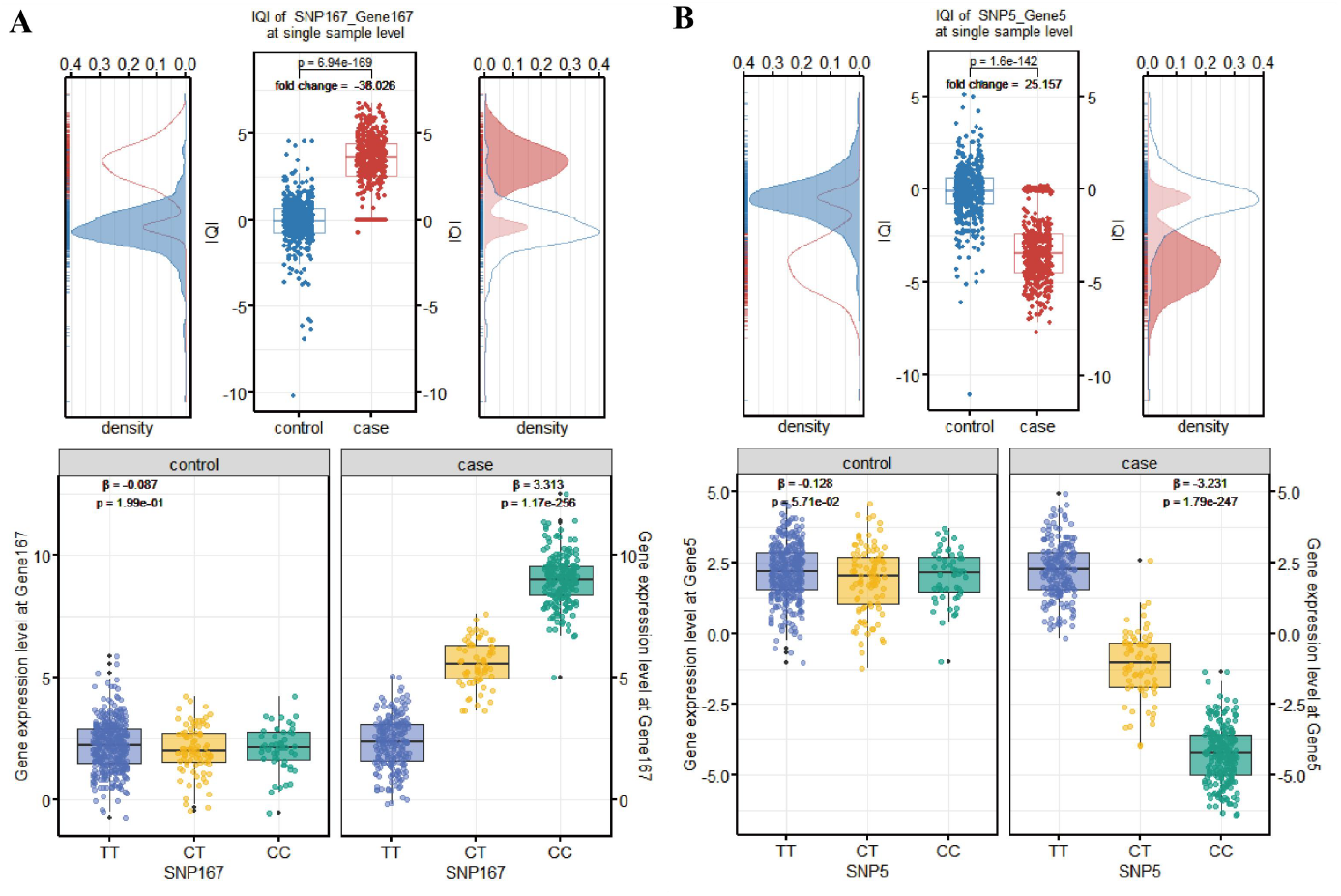
(A) GOE of positive regulation. (B) GOE of negative regulation.

The second type was LOE (Loss of QTL effect) that the QTL regulation only significantly existed in the normal state but not in the disease state (Figure S2). The absolute mean IQI was less than 0.05 in the case but more than 0.05 in the control (the upper half of Figure S2), which is equal to the regression coefficient β (the lower half of Figure S2). The LOE type can be further divided into the loss of positive (Figure S2A) and negative regulation (Figure S2B). In accordance with the IQI, the regulatory effect on SNP to gene expression was positive in the control (p < 0.05, the mean IQI > 0, β > 0) but not significant (p > 0.05) in the case (the lower half of Figure S2A). In Figure S2B, the regulatory effect was not significant in the control (p > 0.05) but negative (p < 0.05, the mean IQI < 0, β < 0) in the case.

**Figure S2.**
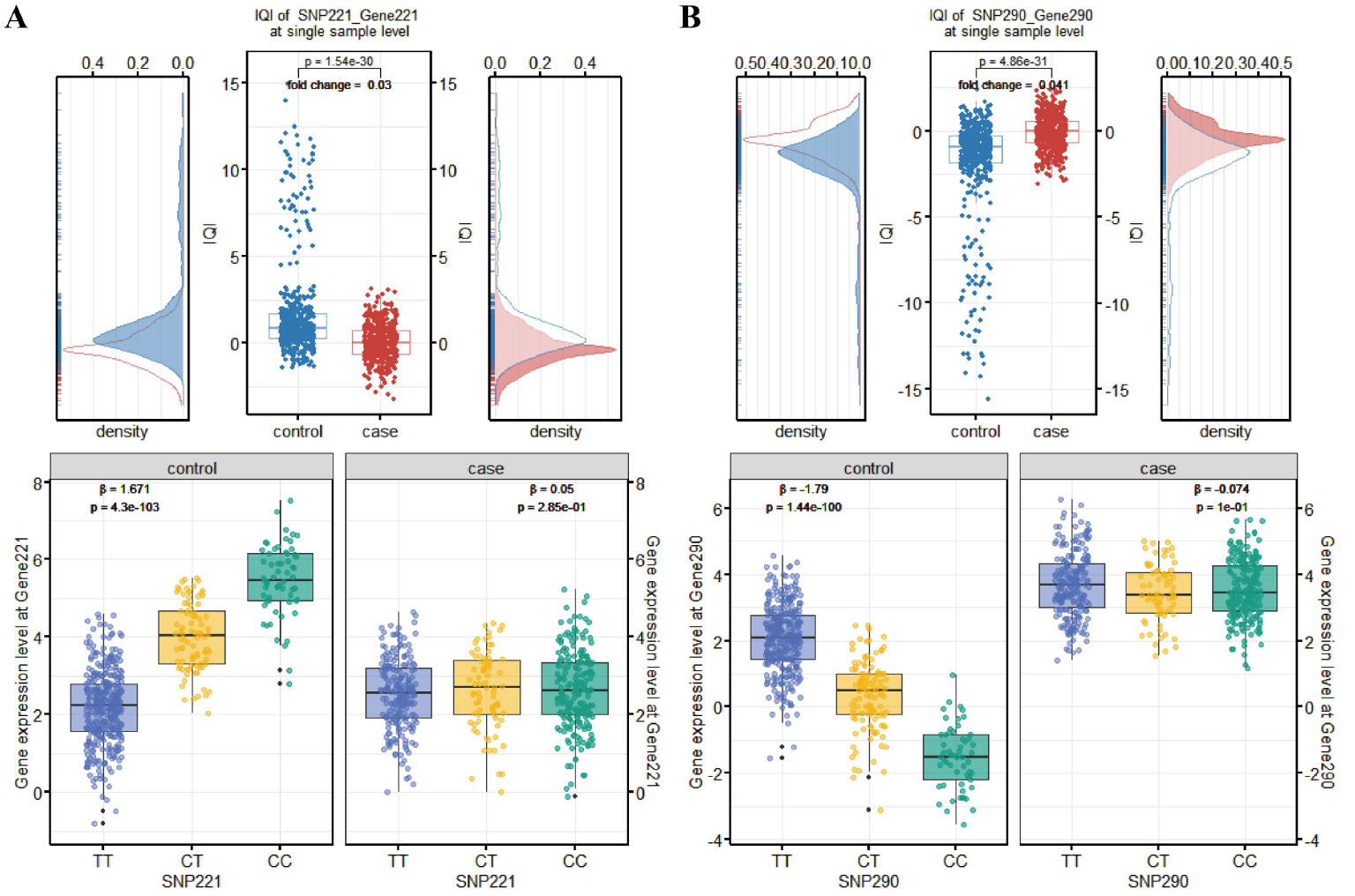
(A) LOE of positive regulation. (B) LOE of negative regulation.

The third type was ROE (Reversal of QTL effect) that the QTL effects significantly exited in the normal and disease state but were in an opposite direction (Figure S3). The absolute mean IQI was both more than 0.05 in the case and control (the upper half of Figure S3), which were equal to the regression coefficient β (the lower half of Figure S3). The ROE type can be further divided into the reversal from positive to negative (Figure S3A) and from negative to positive regulation (Figure S3B). In accordance with the IQI, the regulatory effect on SNP to gene expression was both significant and the regression coefficient β was more than 0 in the control but less than 0 in the case (the mean IQI was more than 0 in the control but less than 0 in the case, the lower half of Figure S3A). In Figure S3B, the regression coefficient β was less than 0 in the control but more than 0 in the case (the mean IQI < 0 in the control but > 0 in the case).

**Figure S3.**
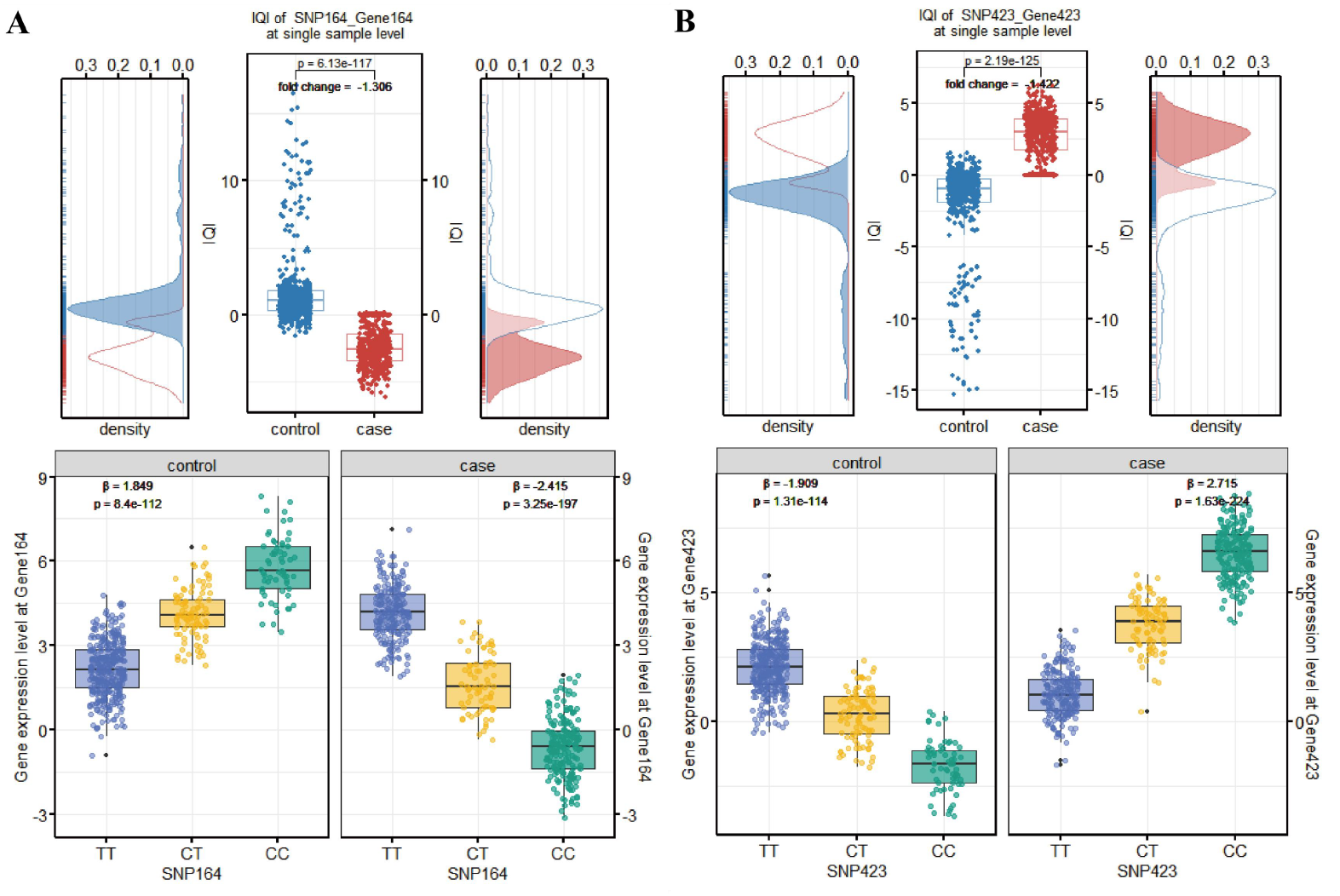
(A) ROE from positive to negative regulation. (B) ROE from negative to positive regulation.

The fourth type was SOE (Strengthening of QTL effect) that the QTL effects significantly exited both in the normal and disease state, but were significantly strengthened in the disease status compared to the normal status. (Figure S4). The absolute mean IQI was both less than 0.05 in the case and control (the upper half of Figure S4) and the control was less than case 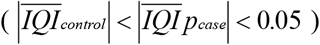. It was equal to the regression coefficient β (the lower half of Figure S4). The SOE type can be further divided into the strengthening of positive (Figure S4A) and negative regulation (Figure S4B). In accordance with the IQI, the absolute regression coefficient β in control was less than case and both were more than 0 (the mean IQI in both case and control were more than 0, the lower half of Figure S4A). In Figure S4B, the regulatory effect was negative and the absolute regression coefficient β in both case and control were less than 0 (the mean IQI in both case and control was less than 0).

**Figure S4.**
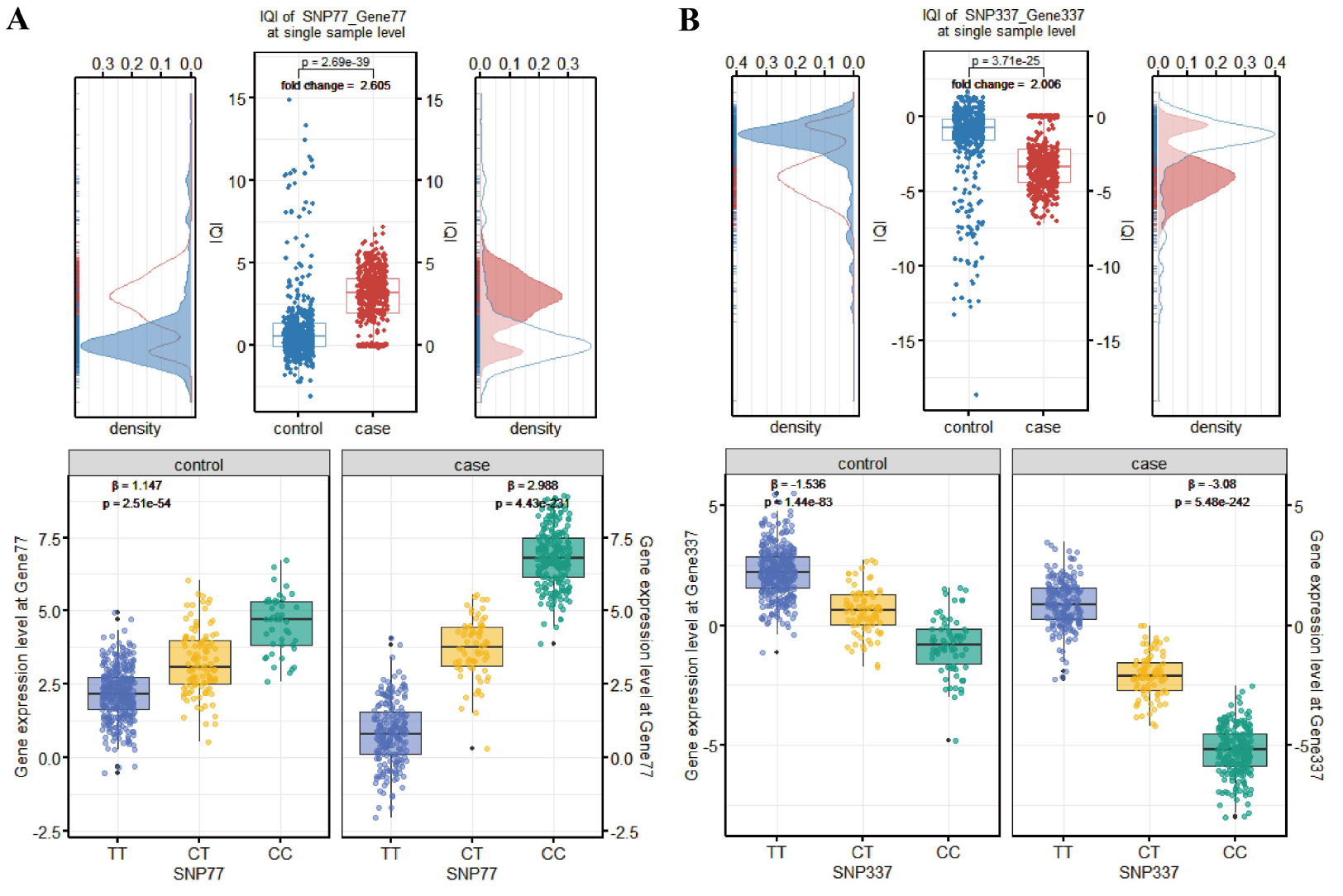
(A) SOE of positive regulation. (B) SOE of negative regulation.

The fifth type was WOE (Weakening of QTL effect) that the QTL effects significantly exited both in the normal and disease state, but were significantly weakened in the disease state compared to the normal state (Figure S5). The absolute mean IQI was both less than 0.05 in the case and control (the upper half of Figure S5) and the control was more than case 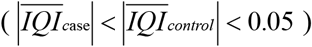. It was equal to the regression coefficient β (the lower half of Figure S5). The WOE type can be further divided into the weakening of positive (Figure S5A) and negative regulation (Figure S5B). In accordance with the IQI, the absolute regression coefficient β in control was more than case and both were more than 0 (the mean IQI in both case and control were more than 0, the lower half of Figure S5A). In Figure S5B, the regulatory effect was negative and the absolute regression coefficient β in both case and control was less than 0 (the mean IQI in both case and control was less than 0).

**Figure S5.**
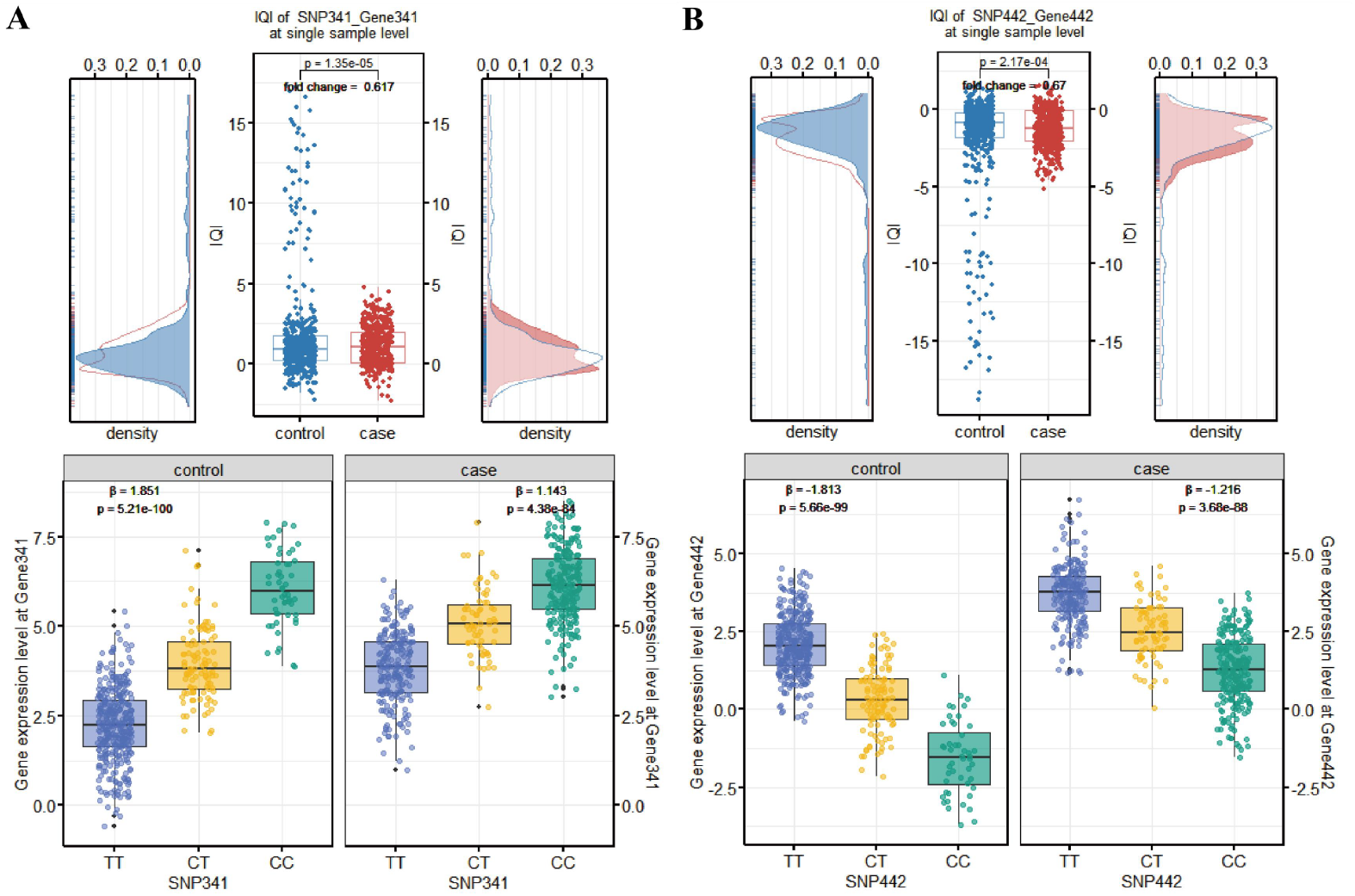
(A) WOE of positive regulation. (B) WOE of negative regulation.

For all the 434 differential QTLs, we calculated the statistical power under different sample size (ranged from 10 to 500) for each group as well as the sample size required to achieve a power of 0.8. More details about all the 434 differential eQTLs can be found in Supplementary Table 1. Notably, the outcomes for the 10 differential QTLs described above (solid lines) and the 10 differential QTLs randomly selected (dashed lines) were displayed in Figure S6. Moreover, the sample size required when power to reach 0.8 and the power for these 20 differential QTLs when the sample size is 500 were shown in Table S2.

**Figure S6.**
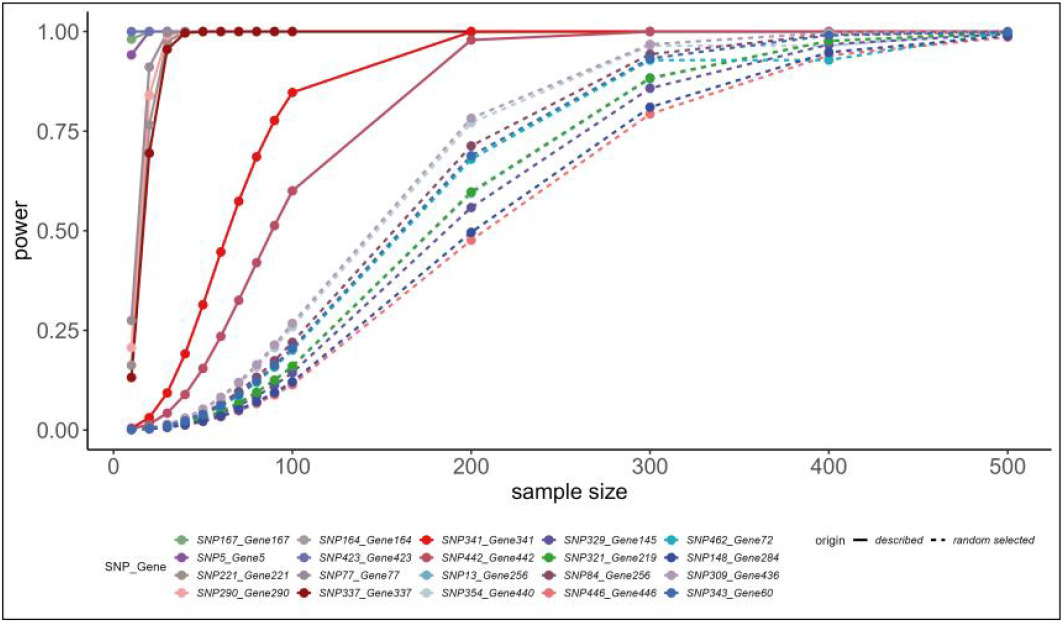
Power under different sample size for the 20 differential QTLs.

**Table S2.**
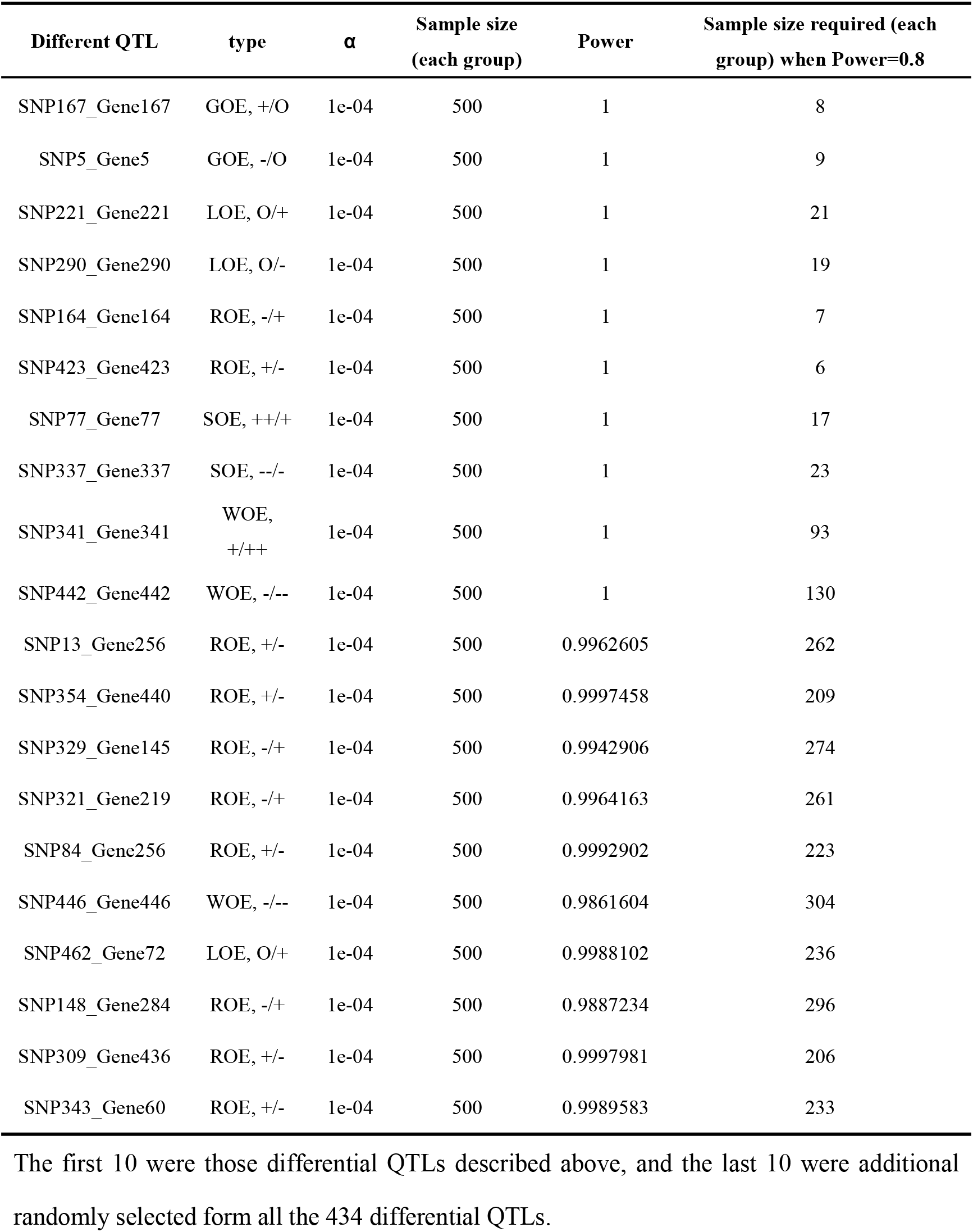
Power for the 20 differential QTLs when sample size=500 and the significance level (α)=1e-04, as well as the sample size required when power=0.8.

### 3. Software for IQI analysis and QTL effect characterization

Here, we first present a hypothesis testing method, named Differential QTL (DQ) analysis framework, to accurately detect the differences between QTL effect in health and disease. DQ analysis framework can be carried out directly by our software, IQI v1.0, which can be freely available at http://www.onethird-lab.com/IQI/. There are four parts in the website, including *Home, Software, Contact us, and Links*. The users can obtain the introduction of IQI v1.0 at *Home*, download the software at *Software*, find our contact information at *Contact us*, and go to relative websites through *Links* (Figure S7).

**Figure S7.**
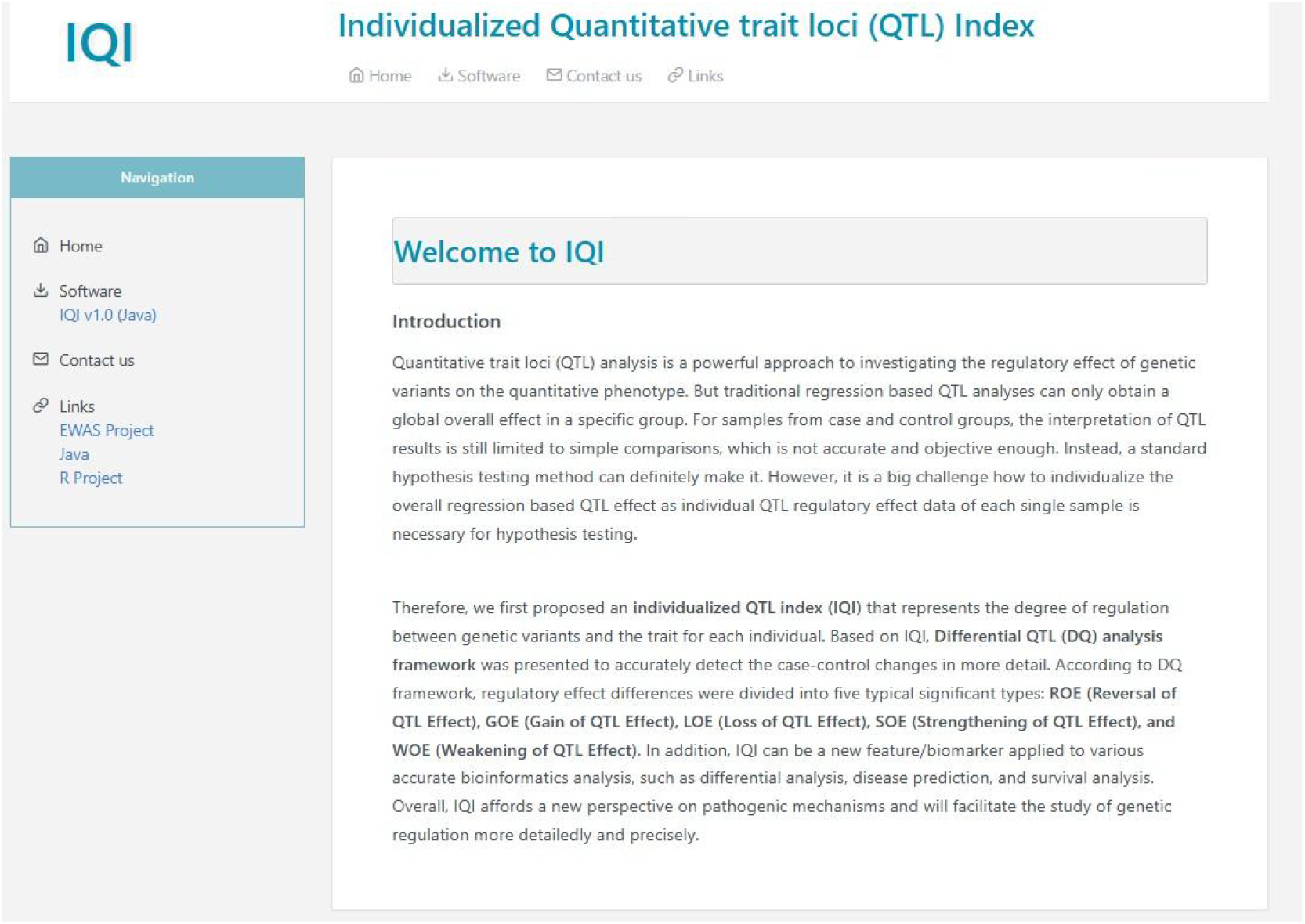
Homepage of the website.

The codes in IQI v1.0 was written by Java which needs to install Java programming environment before use. The IQI v1.0 as a jar package can be downloaded by clicking *IQI v1*.*0 (Java, Command-line running)* at the *Software* (Figure S8). In addition, the introduction of parameters in the command line, input, and output files can be obtained as well.

**Figure S8.**
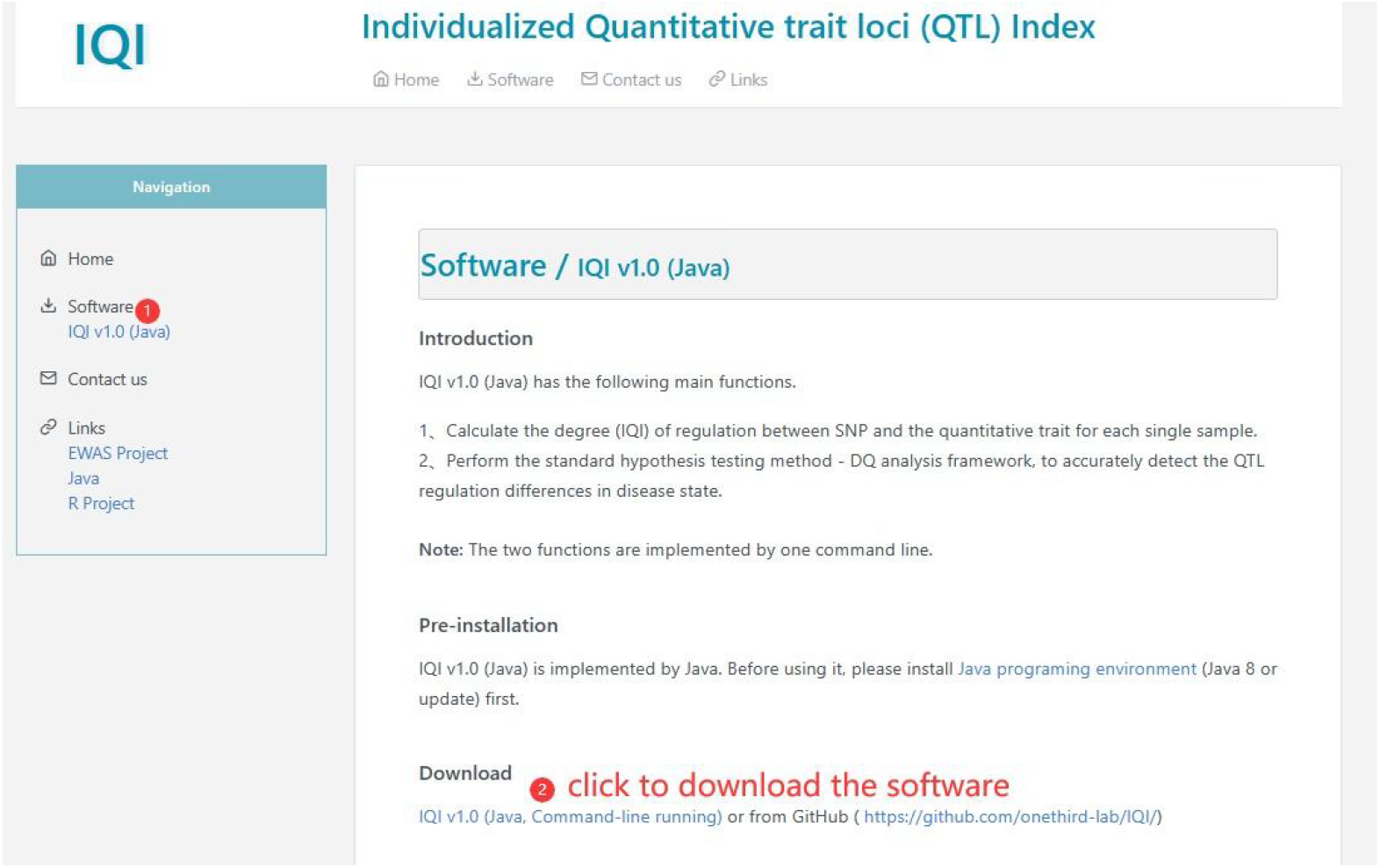
The introduction and usage of IQI v1.0.

IQI v1.0 perform the DQ analysis framework by calculating IQI of the SNP-trait pairs for each single sample between cases and controls first. The software needs four input files, including two trait data and two genotype data of the cases and controls. Three output files can be generated including two files of IQI in the cases and controls and the DQ analysis results. DQ analysis framework can be carried out by a command line, such as:

*java -jar IQI*.*jar -DQ*.*test -input*.*pheno_1 pheno_1*.*txt -input*.*pheno_0 pheno_0*.*txt -input*.*geno_0 SNP_0*.*txt -input*.*geno_1 SNP_1*.*txt -output*.*IQI_0 IQI_0*.*txt -output*.*IQI_1 I QI_1*.*txt -output*.*eQe_test eQe_test*.*txt*

There are eight parameters in the command, including

1. *-DQ*.*test* : the command to perform the DQ analysis framework;
2. *-input*.*pheno_1*: input the trait information of the case group;
3. *-input*.*pheno_0*: input the trait information of the control group;
4. *-input*.*geno_0*: input the SNP genotype data of the case group;
5. *-input*.*geno_1*: Input the SNP genotype data of the control group;
6. *-output*.*IQI_0*: Specify the file name of the IQI values of the SNP-trait pairs of all samples in the control group;
7. *-output*.*IQI_1*: Specify the file name of the IQI values of the SNP-trait pairs of all samples in the case group;
8. *-output*.*eQe_test*: Specify the file name of the results of DQ analysis framework.

The input trait file (*pheno_1*.*txt* and *pheno_0*.*txt*) consists of the name of the quantitative trait and the values of each sample (Table S3) and the SNP genotype file (*SNP_1*.*txt* and *SNP_0*.*txt*) includes the name of SNPs, two alleles, the minor allele and the genotype of each sample (Table S4). Noteworthy, the order of samples in the four input files should be consistent.

**Table S3.** An example of the input trait file. Column 1 is the ID or name of a trait and column 2 to the last is the trait information of each sample.

| Gene expression | Sample<sub>1</sub> | Sample<sub>2</sub> | ... | ... | Sample<sub>n</sub> |
| --- | --- | --- | --- | --- | --- |
| Gene<sub>1</sub> | 7.4095 | 6.9150 |  |  | 6.4904 |
| Gene<sub>2</sub> | 5.2325 | 5.4984 |  |  | 5.2639 |
| ... |  |  |  |  |  |
| ... |  |  |  |  |  |
| Gene<sub>n</sub> | 4.4334 | 5.0982 |  |  | 4.8504 |

**Table S4.** An example of the input SNP genotype file. Column 1 is the name of an SNP, column 2 is the two alleles, column 3 is the minor allele, and column 4 to the last is the genotype of each sample.

| SNP | Allele | Minor.allele | Sample<sub>1</sub> | Sample<sub>2</sub> | ... | ... | Sample<sub>n</sub> |
| --- | --- | --- | --- | --- | --- | --- | --- |
| SNP<sub>1</sub> | G/A | G | GA | AA |  |  | AA |
| SNP<sub>2</sub> | G/T | G | GT | TT |  |  | TT |
| ... |  |  |  |  |  |  |  |
| SNP<sub>n</sub> | T/C | T | CC | CC |  |  | CT |

There are three output files, including the IQI of all samples in the case and control groups (*SNP_1*.*txt* and *SNP_0*.*txt*, Table S5) and the DQ analysis results (*eQe_test*.*txt*) where are eighteen columns:

1. SNP_Gene: the name of SNP-trait pair;
2. change_type: change type of QTL effect;
3. beta_1: the traditional regression based QTL analysis results of the case group;
4. SE_1: the standard error in the case group;
5. p_1: the p value in the case group;
6. beta_0: the traditional regression based QTL analysis results of the control group;
7. SE_0: the standard error in the control group;
8. p_0: the p value in the control group;
9. dx1_x0: the mean difference between case and control groups;
10. Sx1_x0: the standard error between case and control groups;
11. df: the degree of freedom;
12. eQe_mean_1: the average IQI in the case group;
13. eQe_mean_0: the average IQI in the control group;
14. sample_num_1: the sample size of the case group;
15. sample_num_0: the sample size of the control group;
16. fold_change: the fold change value;
17. t: the DQ statistics;
18. tp: the p value.

**Table S5.** An example of the output IQI file. Column 1 is the name of the SNP-trait pair, and column 2 to the last is IQI of each sample.

| **SNP_Gene** | **Sample<sub>1</sub>** | **Sample<sub>2</sub>** | **...** | **...** | **Sample<sub>n</sub>** |
| --- | --- | --- | --- | --- | --- |
| **SNP<sub>1</sub>-Gene<sub>1</sub>** | -0.103258 | 0.349470 |  |  | 0.175838 |
| **SNP<sub>2</sub>-Gene<sub>2</sub>** | 0.054237 | 0.298285 |  |  | 0.150084 |
| ... |  |  |  |  |  |
| ... |  |  |  |  |  |
| **SNP<sub>n</sub>-Gene<sub>n</sub>** | -0.116119 | -0.545830 |  |  | 0.197740 |

### 4. IQI based disease prediction

**Table S6.**
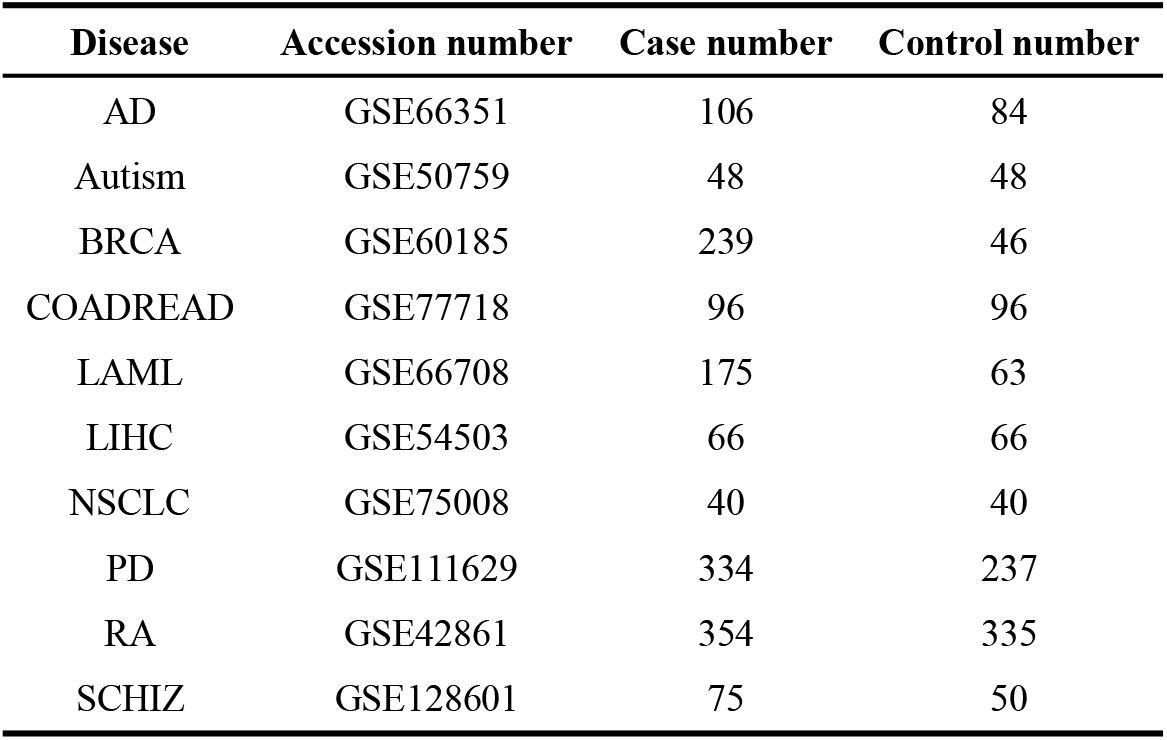
Datasets used for disease prediction analysis.

**Table S7.** Accuracy of SNP, methylation, and IQI and their combined use as separate predictors utilizing SVM model for all 10 diseases.

| disease | SNP | methylation | IQI | SNP and methylation | SNP, methylation and IQI |
| --- | --- | --- | --- | --- | --- |
| RA | 0.528 | 0.727 | 0.895 | 0.727 | 0.917 |
| Autism | 0.648 | 0.669 | 0.990 | 0.660 | 0.953 |
| SCHIZ | 0.599 | 0.670 | 1.000 | 0.736 | 0.998 |
| PD | 0.585 | 0.635 | 0.938 | 0.614 | 0.945 |
| AD | 0.612 | 0.954 | 0.999 | 0.963 | 0.997 |
| BRCA | 0.839 | 0.960 | 0.984 | 0.954 | 0.994 |
| LIHC | 0.621 | 0.952 | 0.954 | 0.946 | 0.978 |
| COADRE | 0.565 | 0.953 | 0.926 | 0.948 | 0.963 |
| NSCLC | 0.613 | 1.000 | 0.950 | 0.991 | 1.000 |
| LAML | 0.735 | 0.996 | 0.981 | 0.993 | 0.993 |

**Table S8.** AUC of SNP, methylation, and IQI and their combined use as separate predictors utilizing SVM model for all 10 diseases.

| disease | SNP | methylation | IQI | SNP and methylation | SNP, methylation and IQI |
| --- | --- | --- | --- | --- | --- |
| RA | 0.523 | 0.727 | 0.894 | 0.726 | 0.916 |
| Autism | 0.642 | 0.670 | 0.990 | 0.662 | 0.953 |
| SCHIZ | 0.500 | 0.622 | 1.000 | 0.694 | 0.998 |
| PD | 0.500 | 0.585 | 0.934 | 0.558 | 0.943 |
| AD | 0.580 | 0.950 | 1.000 | 0.961 | 0.997 |
| BRCA | 0.500 | 0.899 | 0.985 | 0.874 | 0.983 |
| LIHC | 0.618 | 0.953 | 0.953 | 0.946 | 0.978 |
| COADRE | 0.564 | 0.953 | 0.926 | 0.948 | 0.963 |
| NSCLC | 0.613 | 1.000 | 0.950 | 0.991 | 1.000 |
| LAML | 0.500 | 0.992 | 0.975 | 0.989 | 0.990 |

### 5. The DQ statistics obey T distribution

We observed that the DQ statistics obey T distribution when with a small sample size and normal distributions with a large sample size.

Proof: According to the formula for t distribution:

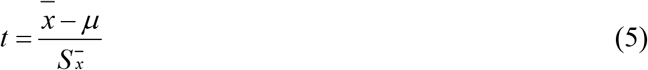

Where 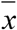 is the sample mean, *μ* is the population mean, as both less than 0.05 in the case astandard deviation. is the sample

Let *x* = 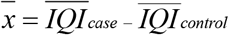 in the Eq (5), i.e.

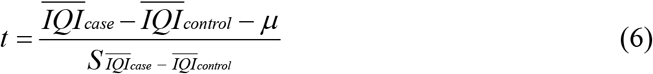

of which,

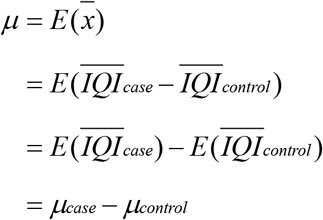

Based on the null hypothesis H_0_ that the QTL effects (mean IQI) are the same between cases and controls, i.e. in the Eq (6), we can observe that

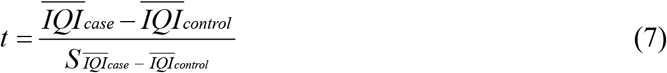

Let *t* = *DQ* in the Eq (7), i.e.

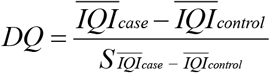

So that, the DQ statistics obey T distribution. When with a large sample size, the degrees of freedom gradually increase and the t distribution gradually approaches the normal distribution.

### 6. IQI based survival analysis

**Table S9.** Datasets used for survival analysis.

| disease | Accession number | sample number |
| --- | --- | --- |
| pancreatic cancer | GSE165764 | 28 |
| breast cancer | GSE39004 | 56 |
| ependymoma | GSE224218 | 141 |
| pediatric low-grade glioma | GSE135017 | 110 |
| pediatric brain tumor | GSE155660 | 28 |
| pediatric adrenocortical tumors | GSE179175 | 53 |
| neuroendocrine tumours | GSE211483 | 40 |

**Table S10.**
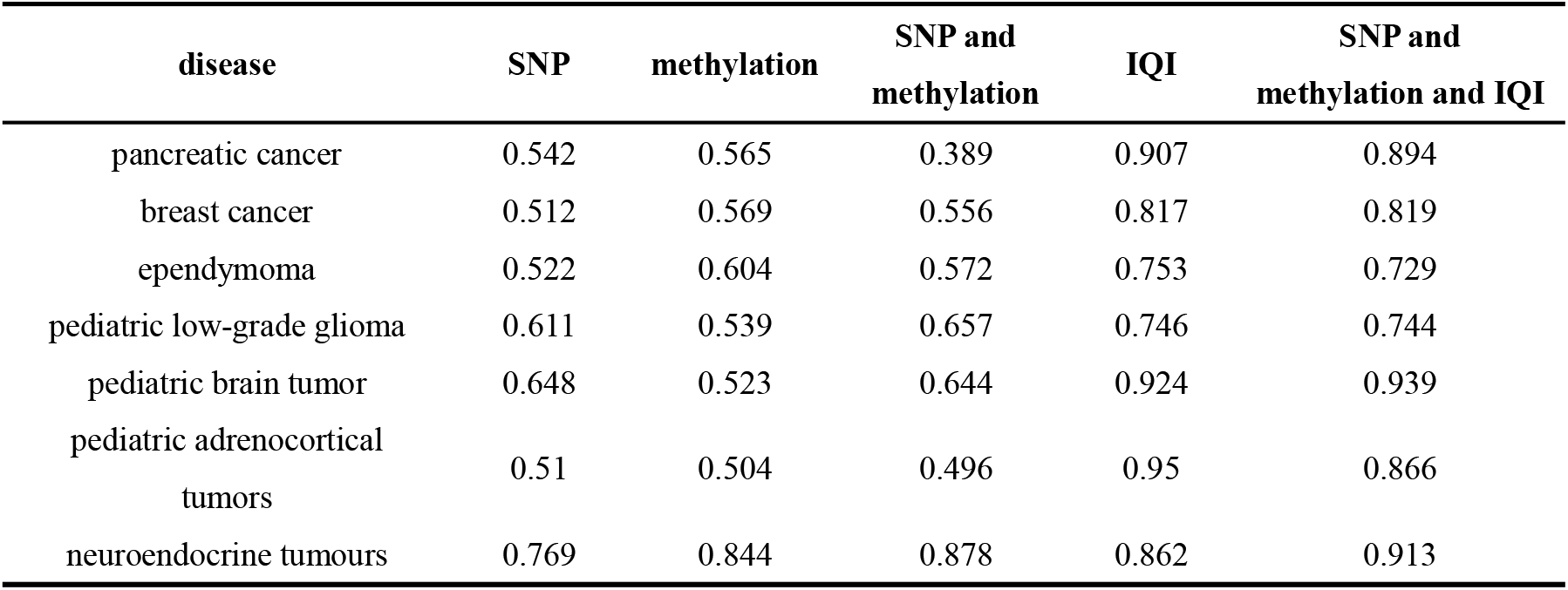
The C-index of IQI, SNP, and methylation in prognosis analysis.

### 7. The predictive performance comparisons of IQI before and after data shuffling

**Figure S9.**
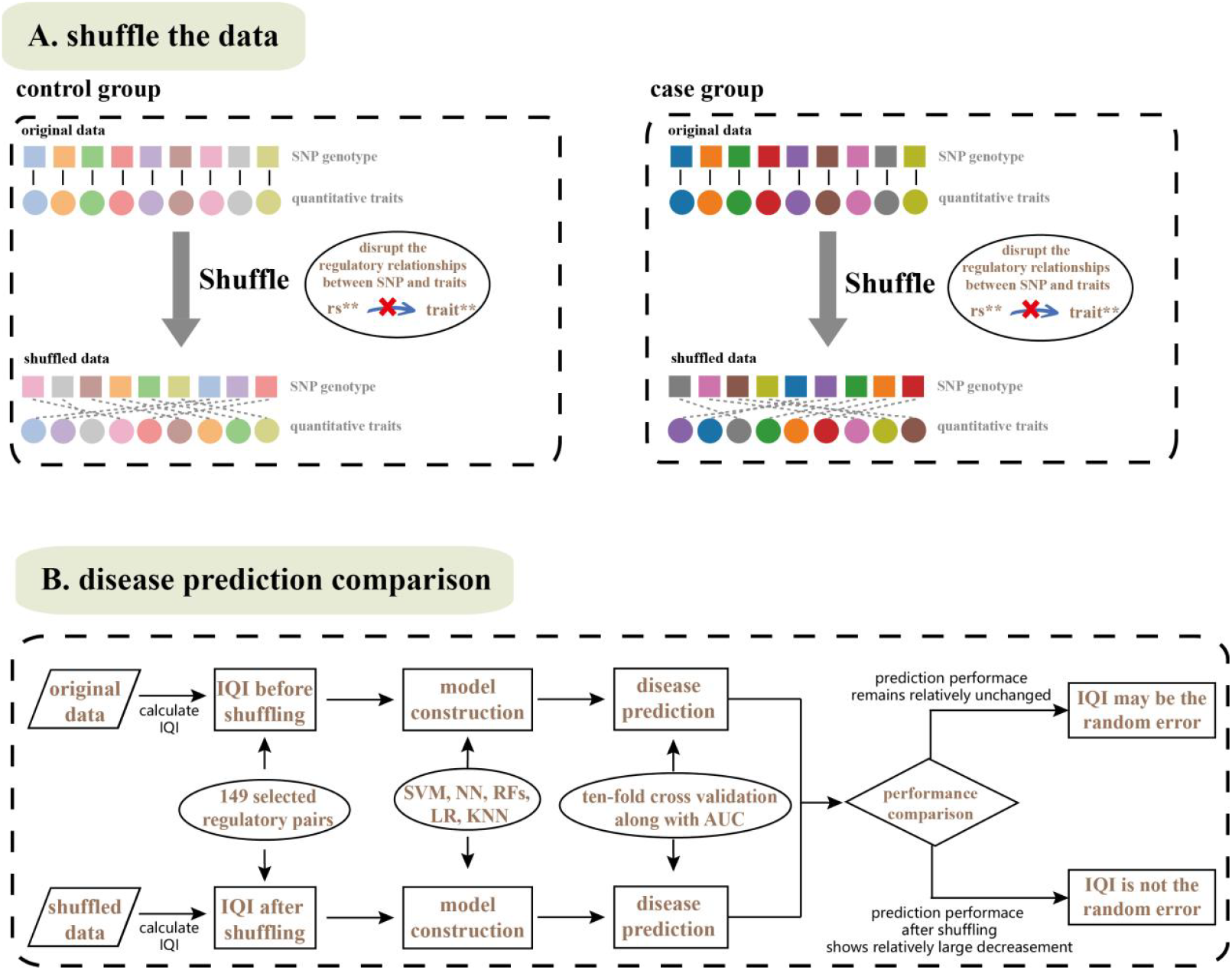
The schematic diagram of the shuffling test.

**Figure S10.**
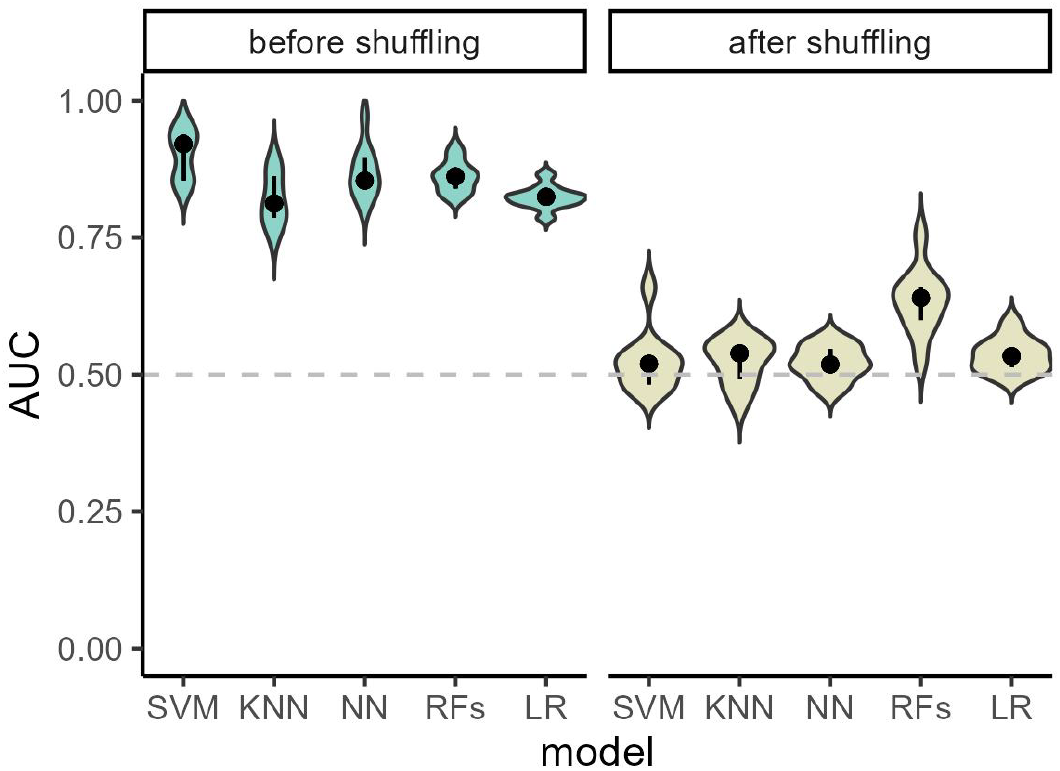
The AUC of ten-fold cross validation for KNN, LR, NN, RFs, and SVM before and after shuffling.

